# Biophysical modeling of membrane curvature generation and curvature sensing by the glycocalyx

**DOI:** 10.1101/2024.09.07.611813

**Authors:** K. Xiao, S. Park, J. C. Stachowiak, P. Rangamani

**Affiliations:** Department of Mechanical and Aerospace Engineering, University of California San Diego, La Jolla CA 92093, USA; Department of Biomedical Engineering, University of Texas at Austin, Austin, TX 78712, USA; Department of Pharmacology, School of Medicine, University of California San Diego, La Jolla CA 92093, USA

**Keywords:** Glycocalyx, membrane bending, curvature generation, curvature sensing, Helfrich energy, polymer brush theory

## Abstract

Generation of membrane curvature is fundamental to cellular function. Recent studies have established that the glycocalyx, a sugar-rich polymer layer at the cell surface, can generate membrane curvature. While there have been some theoretical efforts to understand the interplay between the glycocalyx and membrane bending, there remain open questions about how the properties of the glycocalyx affect membrane bending. For example, the relationship between membrane curvature and the density of glycosylated proteins on its surface remains unclear. In this work, we use polymer brush theory to develop a detailed biophysical model of the energetic interactions of the glycocalyx with the membrane. Using this model, we identify the conditions under which the glycocalyx can both generate and sense curvature. Our model predicts that the extent of membrane curvature generated depends on the grafting density of the glycocalyx and the length of the polymers constituting the glycocalyx. Furthermore, when coupled with the intrinsic membrane properties such as spontaneous curvature and a line tension along the membrane, the curvature generation properties of the glycocalyx are enhanced. These predictions were tested experimentally by examining the propensity of glycosylated transmembrane proteins to drive the assembly of highly-curved filopodial protrusions at the plasma membrane of adherent mammalian cells. Our model also predicts that the glycocalyx has curvature sensing capabilities, in agreement with the results of our experiments. Thus, our study develops a quantitative framework for mapping the properties of the glycocalyx to the curvature generation capability of the membrane.

**Significance Statement:** The glycocalyx is a dense layer of glycosylated transmembrane proteins and lipids distributed on the extracellular surface of eukaryotic cells. It is known to mediate cell-cell interactions and protect cells from invasion by pathogens. However, recently it has been found to play a role in generating membrane curvature, which is essential to diverse cellular functions spanning from endocytosis to cell division. Experiments have revealed that different membrane shapes: spherical vesicles, pearls, and tubes are regulated by the glycocalyx. However, currently, we lack a quantitative physical explanation of how glycocalyx properties determine membrane geometries. Here we develop a polymer brush theory-based model, which suggests that the interplay between glycocalyx polymers and membrane bending captures the wide variety of membrane shapes from spherical buds to elongated pearl-like shapes found in previously published experiments. We predict that the physical properties of glycocalyx polymers, line tension, and membrane elastic parameters play significant roles in regulating membrane morphologies. We find that the glycocalyx prefers higher local membrane curvature indicating that the glycocalyx can sense local curvature.

## 1 Introduction

Cellular membranes are capable of bending into highly curved structures and such membrane curvature generation is critical for cellular function (*1–4*). From a biophysical standpoint, commonly studied mechanisms for curvature generation include protein insertion, scaffolding, and crowding (*5–11*). In addition to membrane insertions, many studies have shown that different polymer structures including the glycocalyx play an important role in regulating membrane morphologies (*12–34*). The mechanism of membrane bending in the presence of polymers seems to stem from the crowding-based steric forces between the polymer chains (*6, 7, 33–35*). Experimental studies by Shurer et al. (*30*) have shown that bulky brush-like glycocalyx polymers are sufficient to induce a variety of curved membrane features (Fig. 1A), including flat membranes, spherical membrane blebs, and unduloids, in a polymer density-dependent manner. These findings collectively indicate that the glycocalyx is a potent driver of membrane deformation and affects the membrane reshaping processes. The glycocalyx is a thick sugar-rich layer of proteins and carbohydrates concentrated on most cell surfaces in a complex structure (*36–38*). These molecules have bottlebrush-like molecular structures formed by polymer backbones and attached sugar sidechains (Fig. 1B).

**Figure 1:**
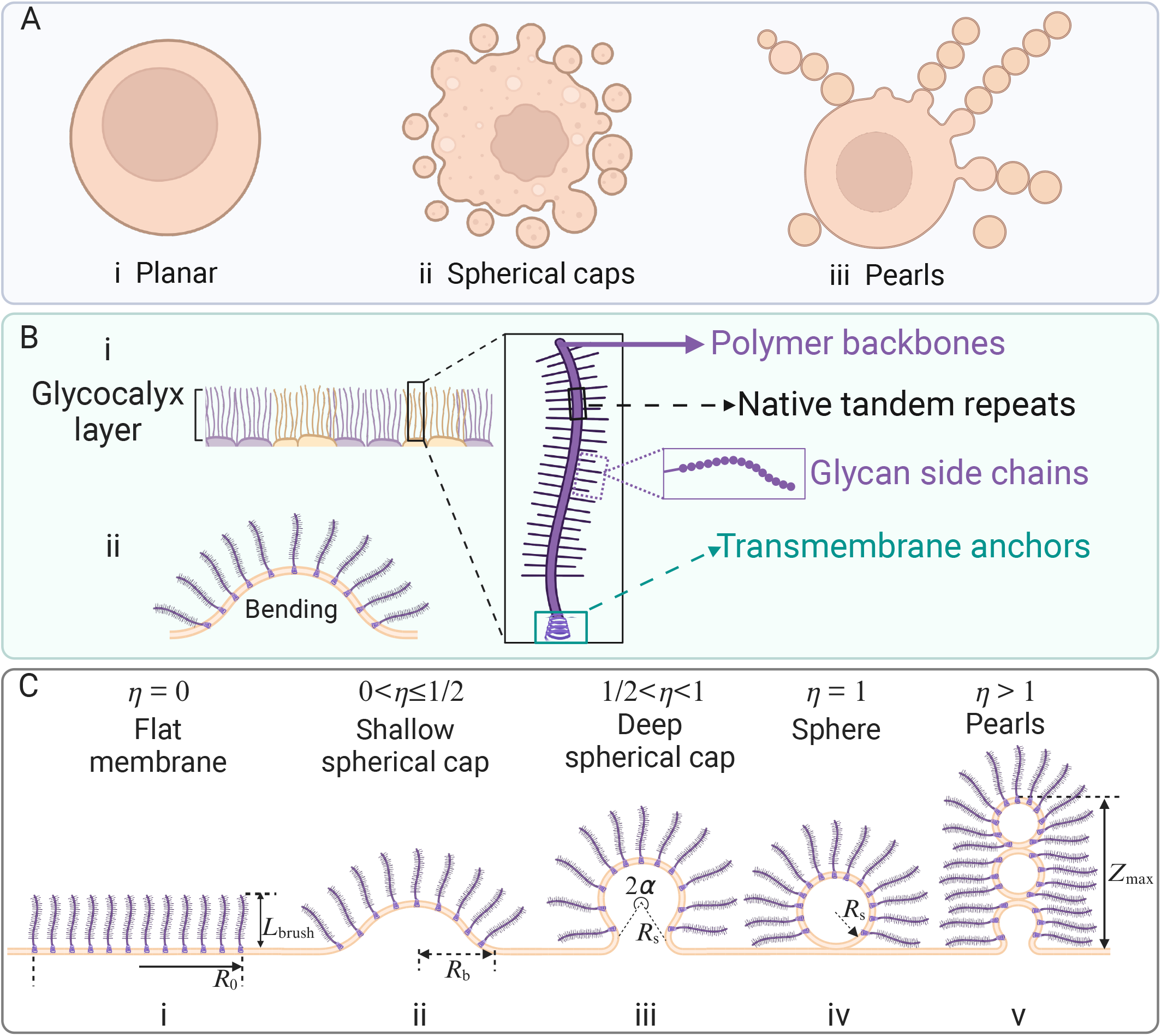
Polymers such as the glycocalyx can induce a variety of membrane curvatures. (A) Curvatures generated by the glycocalyx can be described as (i) mostly flat, (ii) spherical buds, and (iii) pearled shapes. (B) Schematic showing the polymer structure of glycocalyx constituents such as mucins. We focus on the brush-like regime. At high polymer density, the stretched polymer chains leads to a decrease of the chain entropy and a related increase of the chain free energy. To reduce the entropy cost of the system, bending of the membrane provides more space and gives more orientational degrees of freedom for the polymer chains. (C) Progression of membrane shape as a function of shape parameter *η*. The key geometric variables are shown.

Biophysical modeling enables a quantitative understanding of how polymer networks such as glycocalyx can induce membrane curvature. The shape remodeling of membrane caused by anchored polymers has been investigated in detail by an extensive body of theoretical studies (*39–47*). Analytical methods (*39–42*) and scaling theory (*43*) have been used to demonstrate that the entropic pressure exerted by polymers results in the changes to the shape, bending moduli, and spontaneous curvature of the membrane. Studies using the Canham-Helfrich model and phase-field simulations (*44, 45*) have shown that the formation of membrane tubes and curvature-driven pearling instabilities in membranes can be attributed to the concentration gradient of anchored polymers and the concentration of homogeneous anchored amphiphilic polymers, respectively. Additionally, studies using Monte Carlo simulations (*47*) have shown that polymer-anchored membranes with high tension exhibit lower curvature and that reducing membrane tension can lower the threshold of polymer density for driving curvature. In particular, a crowded layer of polymers on one surface of the membrane will drive membrane curvature whenever the steric forces generated by the polymers are not matched by equivalent forces on the opposite membrane surface. These results suggest that the membrane curvature depends on both the crowding of the biopolymers and on membrane properties. Recently, Shurer et al. (*30*) proposed a theoretical framework by extending fundamental concepts in polymer physics to the glycocalyx. They mapped the effects of glycopolymer density and length onto membrane spontaneous curvature in distinct regimes (mushroom regime and brush regime) and on generating distinct membrane morphologies.

In many of the prior studies, the effective curvature generation capability of the polymers is mapped onto the spontaneous curvature of the membrane, a lumped free parameter in the Helfrich model (*39–41, 43, 45*). In this work, we sought to separate the contributions of the membrane spontaneous curvature and the glycocalyx, we treat the membrane spontaneous curvature as a contribution arising from the internal asymmetries of lipids and transmembrane proteins (*48*). Since the glycocalyx layer is significantly thicker than the membrane thickness, we consider the energy of this polymer layer in terms of its physical characteristics (*49*) and we treat it as a separate contributor to the total energy of the system. Thus, we build an energetic framework that allows us to tune the properties of the glycocalyx and the membrane independently. Studies also have shown that most proteins capable of generating curvature can also sense curvature (*50*), where sensing refers the favorably partitioning of proteins to curved membrane regions. While there has been an extensive body of work on different curvature-sensing proteins, (*51–56*), whether the glycocalyx itself can sense curvature remains unclear. Specifically, if glycosylated proteins partition favorably to regions of higher membrane curvature, then the local density of the glycocalyx may vary with membrane curvature (*57*).

Given the various factors that can alter in cells, such as the density and length of glycocalyx constituent molecules, as well as membrane properties like bending modulus, membrane tension, line tension, and lipid-induced spontaneous curvature, we aim to identify the regimes in which membrane properties can collaborate or compete with the stresses induced by the glycocalyx. Specifically, our model predicts that the density and length of the glycocalyx are important determinants of curvature generation even when there is no intrinsic membrane spontaneous curvature. These predictions were tested in experiments by expressing transmembrane proteins with O-glycosylated mucin domain of varying length and quantifying the resulting increase in the density of positively curved filopodial protrusions at the plasma membrane of adherent mammalian cells. We further show that the formation of buds and pearled shapes on the membrane depends on both the glycocalyx and the membrane properties.

Extending our model to spatial heterogeneities in the distribution of the glycocalyx predicts that regions of high curvature have a higher density of the glycocalyx suggesting that the glycocalyx can sense curvature. These predictions were tested by measuring the relative partitioning of glycosylated transmembrane domains between highly curved cellular protrusions and the surrounding plasma membrane of adherent cells. Thus, our study captures both the curvature-generating and curvature-sensing capabilities of the glycocalyx.

### Model Development

In this work, we develop a biophysical model for the glycocalyx-membrane composite (Fig. 1C). The lipid bilayer is modeled as thin elastic shell which resists out-of-plane bending and we assume that the Helfrich energy is sufficient to capture membrane bending. The glycocalyx is modeled as a polymer layer of height *L*_brush_. The energy of the polymer layer depends on the polymer properties including the grafting density, the length of the polymer, and the interactions between the individual polymer chains and also on the curvature of the underlying substrate (*49*). Our approach is to find the shape that corresponds to the minimal energy state for the combined energy of the membrane and the polymer for spherical deformations. We briefly summarize the model here and include additional details in the supplementary material.

#### Model Assumptions

We assume that the glycocalyx can be represented as a thick layer of brush-like polymers that are anchored to the membrane surface. To build our model, we assume that the glycocalyx is electrically neutral and we do not consider the effect of co- and counterions on the polymer chains and therefore, do not consider the electrostatic effects within a polymer chain. We restrict the shape of glycocalyx polymer anchored membrane patch to a family of spherical caps. This assumption allows us to analytically express the free energy as a function of membrane shape. In addition, we also assume that the membrane is always at mechanical equilibrium and neglect both fluctuations and inertia. The bending modulus, membrane tension, line tension, spontaneous curvature, and area over which the glycocalyx polymer is grafted are set as constant unless otherwise specified.

#### Curvature Generation Model

We consider a system consisting of a cell membrane and a layer of densely anchored glycocalyx polymers above it with the total energy given by a sum of three terms: the energy associated with the cell membrane deformation (*F*_membrane_), the energy contribution associated with the glycocalyx (*F*_glycocalyx_), and the energy associated with the line tension at the interface of glycocalyx-rich and glycocalyx-poor phase (*F*_line tension_)

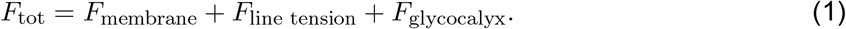

To describe the energy cost of bending the membrane away from its preferred shape, the elastic energy of the membrane is described by the well-known Canham-Helfrich Hamiltonian (*48*) which includes the bending energy of the membrane and the surface tension energy and is given by

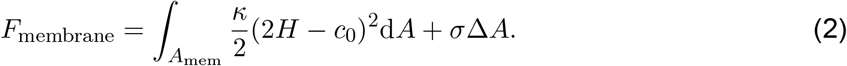

Here κ is the membrane bending rigidity, *H* = (*c*_1_ + *c*_2_)*/*2 is the mean curvature in which *c*_1_ and *c*_2_ are the two principal curvatures, *c*_0_ is the spontaneous curvature, *σ* is the membrane tension, and Δ*A* is the decrease in the in-plane area due to the membrane shape deformation. Here *c*_0_ is the spontaneous curvature which is restricted to asymmetries in the bilayer.

The second term in Eq. (1) is the energy associated with the line tension and is given by

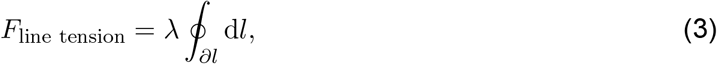

where *λ* represents the strength of line tension along the glycocalyx polymers anchored domain boundary and the in-plane spherical cap base radius. The integral is over the periphery line d*l* of the membrane patch on which the glycocalyx is anchored. We note that the line tension is used in our model is a simplification of the complexities of the heterogeneous composition of the glycocalyx on the cellular membrane.

The last term in Eq. (1) is the energy contribution from glycocalyx polymers which is composed of two terms: the elastic energy, 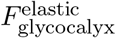, of the polymer brush and the excluded volume interaction of monomers, 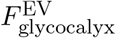, in the polymer brush. Based on Ref. (*49*), the sum of the energy density of these two terms for a single polymer is given by

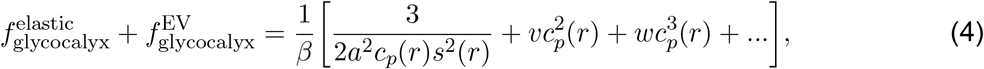

where the first term on the right-hand side describes the elastic stretching of the chains and the last two terms capture the interactions between monomers in the brush. Here, *β* = 1*/*(*k*_*B*_*T*) with *k*_*B*_*T* being the thermal energy, *a* is the monomer size (Kuhn length), *c*_*p*_(*r*) is the local monomer density profile along the thickness, *r* is the radial distance defined from the center of the spherical surface, *s*(*r*) is the area per chain at distance from the polymer grafting surface, and *v* and *w* are the second and third virial coefficients respectively. Integrating this energy density from *R* to *R* + *L*_brush_ and then multiplying the total number of grafted polymers leads to the total energy contribution from glycocalyx polymers (see SI 1.4. Energy contribution of glycocalyx polymer). Note that only the pairwise monomer-monomer interactions with second virial coefficient *v* is considered, and that the ternary interactions with third virial coefficient *w* is neglected.

To minimize the energy, we fixed the area of the membrane on which the glycocalyx is grafted as 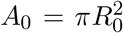, where *R*_0_ is the in-plane radius of the flattened membrane (Fig. 1C). In the brush regime, the grafting density (polymers per unit area) is given by *ρ* = *ξ*^−2^ where *ξ* is the grafting distance and is less than the radius of gyration *R*_*g*_ (*30, 58*). As noted in the assumptions, the membrane deformation can be described by a spherical cap of radius *R*_*s*_ and angle 2*α* (Fig. 1C). This restricted family of shapes can be conveniently specified by a shape parameter *η*, which is defined as the fraction of the surface of spheres of uniform radius *R*_*s*_ covered by the glycocalyx polymers (see SI 1.1). We briefly present the total energy of the glycocalyx-membrane system here,

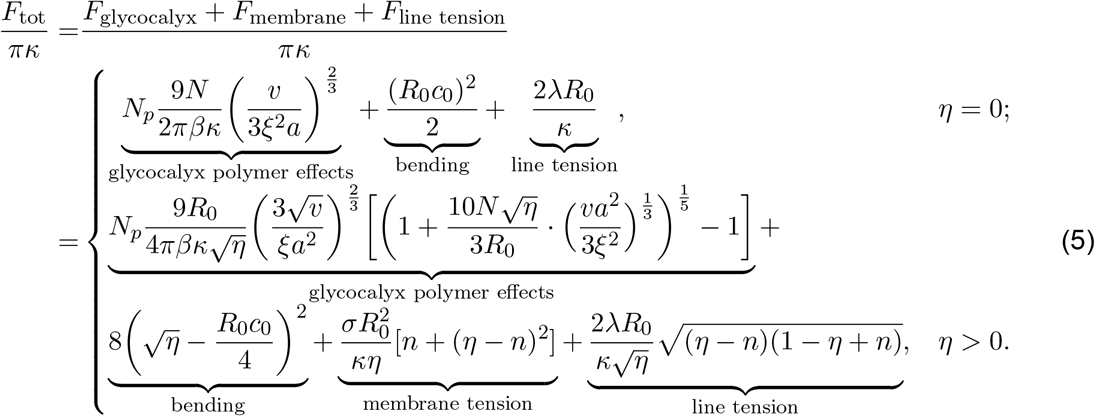

a complete derivation can be found in the Supplementary Material (SI 1). Here *N*_*p*_ is the number of polymer chains grafted on the membrane related to the grafting density as *N*_*p*_ = *ρA*_0_, *N* is the number of monomers in a polymer brush, and *n* = [*η*] is the number of complete unduloids, the brackets denoting the integer part operator. In this work we use the number of monomers, *N*, to capture the length of the polymer due to that the thickness of polymer brush is related to the total number of monomers *N* (SI 1.4).

#### Curvature Sensing by the Glycocalyx

The curvature generation model described above focuses on a single patch of the membrane decorated by the glycocalyx. To quantify the heterogeneous distribution of the glycocalyx on cellular membranes and to estimate its curvature sensing capabilities, we developed a thermodynamic model. We used this model to predict the area fraction of the glycocalyx, *ϕ*, as a function of membrane curvature (*56, 59–62*). The free energy of this curvature sensing model can be written as

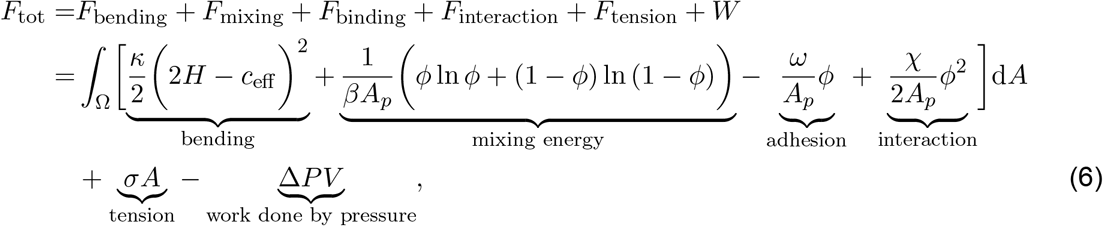

where *c*_eff_ is the effective spontaneous curvature calculated from the curvature generation capabilities of the glycocalyx, *A*_*p*_ is the surface area occupied by the glycocalyx anchor, *ϕ* is the area fraction of glycocalyx on the cell surface, *ω* is the chemical potential of the glycocalyx attachment to the membrane, χ is the measure of the strength of inter-glycopolymer interactions, and Δ*P* is the hydrostatic pressure difference between inside and outside the cell. By minimizing the total free energy in Eq. (6), one can obtain the glycocalyx area fraction *ϕ* on the cell surface and further quantify the curvature sensing ability of glycocalyx. A detailed description of the curvature sensing model of glycocalyx is presented in SI 2. Curvature Sensing Model.

#### Numerical implementation

We determine the global minimum energy state by numerically calculating the total energy of the system for different parameters. For the curvature generating model, we constructed the total free energy of the membrane patch with fixed area *A*_0_ (Eq. (5)). Minimization of this energy as a function of shape factor *η* yields the optimal *η*_min_, which is then used to determine the morphology of the membrane patch. For the curvature sensing model, we numerically solve the equation (S36) derived in the SI 2 for a set of given parameters to obtain the area fraction of glycocalyx on the cell surface. The values of the parameters used in the model are summarized in Tables 1. The code is available on [https://github.com/RangamaniLabUCSD/Glycocalyx-Membrane].

**Table 1:** Model parameter values.

| Parameter | Meaning | Estimated range | References | Value used if not varied |
| --- | --- | --- | --- | --- |
| $\kappa$ | Bending rigidity | $10 \sim 400 k_B T$ | (63–66) | $25 k_B T$ |
| $\sigma$ | Membrane tension | $10^{-4} \sim 1$ pN/nm | (64, 67, 68) | $0.012 k_B T/\text{nm}^2$ |
| $\lambda$ | Line tension | $0 \sim 100$ pN | (64, 69, 70) | $0.5 k_B T/\text{nm}$ |
| $R_0$ | Patch radius | $20 \sim 100$ nm | (47, 64) | 50 nm |
| $\xi$ | Grafting distance | $10 \sim 100$ nm | (30, 71) | 15 nm |
| $a$ | Monomer size in the glycocalyx | $10 \sim 20$ nm | (30, 36, 47, 72–74) | 10 nm |
| $N$ | Number of monomers in a brush | $10 \sim 50$ | (30, 36, 47, 74) | 20 |
| $c_0$ | Spontaneous curvature | $0 \sim 0.1 \text{ nm}^{-1}$ | (30) | $0.04 \text{ nm}^{-1}$ |

## Results

### Glycocalyx alone can regulate membrane bending

We begin by investigating the effects of glycocalyx on the accessible equilibrium configurations of the system as a function of the shape parameter *η*. Recall that *η* characterizes the intermediate shape of the membrane (Fig. 1C).

We first sought to investigate whether the glycocalyx can bend the cell membrane by itself, without any assistance from spontaneous curvature or line tension. We calculated the total free energy as a function of the shape parameter *η* at two different main glycocalyx parameters: the grafting density and the length of the glycocalyx for fixed values of *κ* = 10 *k*_*B*_*T*. For small values of the grafting density or glycopolymer length, our results show that the global minimum of the free energy corresponds to a spherical cap shape (0 *< η <* 1) (Figs. 2A and 2 B). These shapes indicate that the glycocalyx is indeed capable of generating membrane curvature. As we increase the grafting density or the glycopolymer length, the global stable state shifts to a shape corresponding to *η >* 1 (black curve in Fig. 2A and purple curve in Fig. 2B). This result indicates that the glycocalyx is able to bend the membrane into a complete spherical bud without the assistance of line tension and membrane spontaneous curvature.

**Figure 2:**
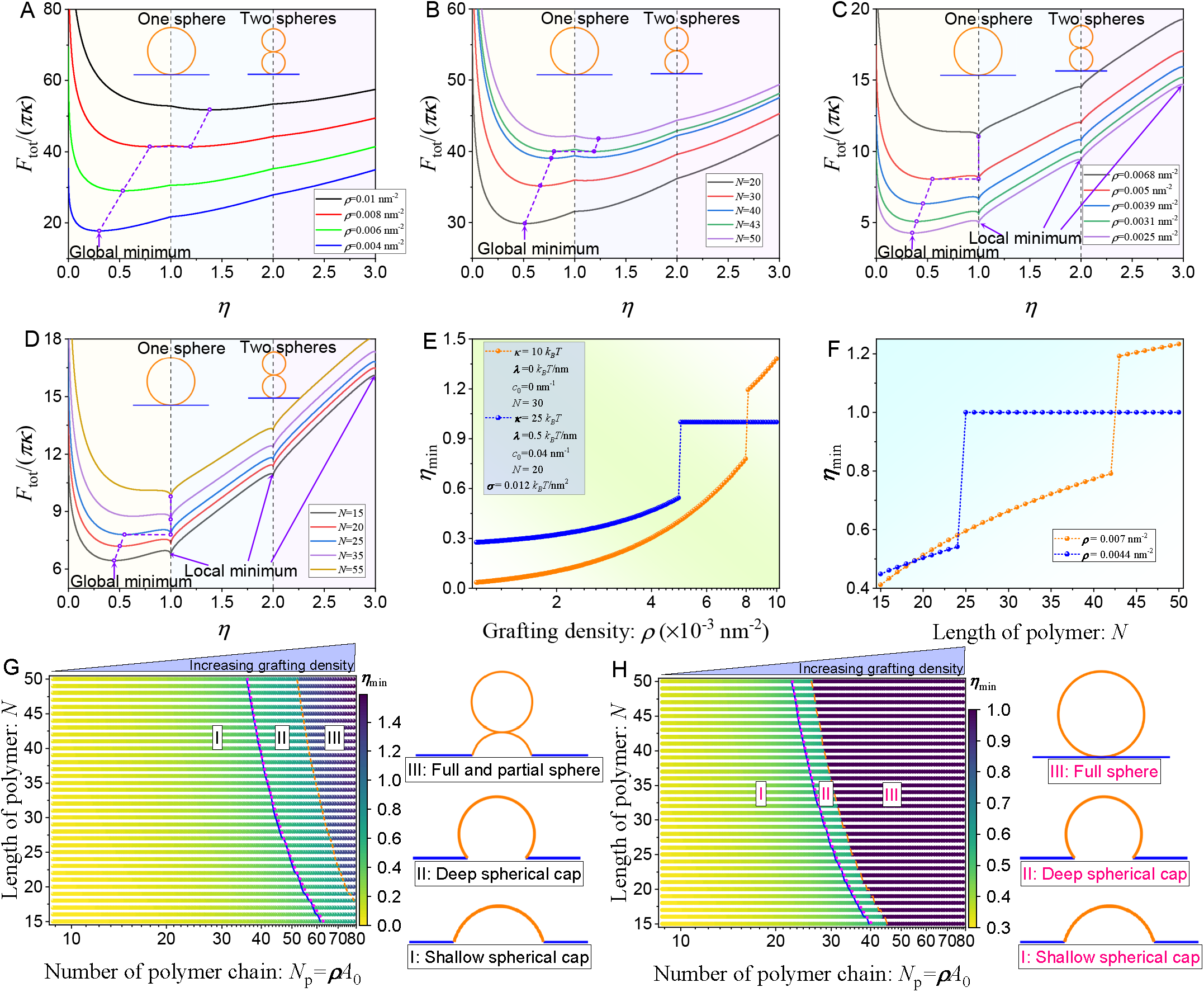
Glycocalyx alone can regulate membrane bending. Total energy profiles as a function of shape parameter *η* for different glycopolymer (A) grafting density *ρ*, and (B) length as a set of parameters are fixed at *κ* = 10 *k*_*B*_*T, λ* = 0 *k*_*B*_*T/*nm, *c*_0_ = 0 nm^−1^, and *N* = 30. Total energy profiles as a function of shape parameter *η* for different glycopolymer (C) grafting density *ρ*, and (D) length as a set of parameters are fixed at *κ* = 25 *k*_*B*_*T, λ* = 0.5 *k*_*B*_*T/*nm, *c*_0_ = 0.04 nm^−1^, and *N* = 20. The dependence of minimum shape parameter *η*_min_ on glycopolymer (E) grafting density and (F) length. Phase diagrams present the optimal shape parameter *η*_min_ as a function of the number of polymer chain, *N*_*p*_, and the length of polymer, *N*, as a set of parameters are fixed at (G) *κ* = 10 *k*_*B*_*T, λ* = 0 *k*_*B*_*T/*nm, *c*_0_ = 0 nm^−1^, and *N* = 30; and (H) *κ* = 25 *k*_*B*_*T, λ* = 0.5 *k*_*B*_*T/*nm, *c*_0_ = 0.04 nm^−1^, and *N* = 20.

We next investigated how this global minimum energy state is affected by increasing the bending rigidity of the membrane. We found that when the membrane bending rigidity increases, a finite value of line tension and membrane-associated spontaneous curvature is needed to promote bud formation (Figs. 2C and 2D). For *κ* = 25 *k*_*B*_*T*, line tension *λ* = 0.5 *k*_*B*_*T/*nm, and spontaneous curvature *c*_0_ = 0.04 nm^−1^, we found that the energy curves have multiple local minima separated by energy barriers. At low glycopolymer grafting density or short glycopolymer length, the global minimum of the free energy corresponds to a spherical cap shape (0 *< η <* 1), while other local minimum states corresponding to one sphere and a stack of spheres (purple, green, and blue curves in Fig. 2C; black and red curves in Fig. 2D). In contrast, at high grafting density or long glycopolymer length, the global free energy minimum is shifted to a spherical shape. In contrast, the spherical cap state is now a local minimum and also will disappear as the grafting density or glycopolymer length exceeds a threshold (black curve in Fig. 2C; and brown curve in Fig. 2D). To better visualize the effect of glycopolymer grafting density and length on the shift in globally stable shape, we plot the optimal shape parameter *η*_min_ as a function of the grafting density and glycopolymer length (Figs. 2E and 2F). When the grafting density or glycopolymer length increases, the optimal shape parameter shifts gradually and then jumps to 1 (blue curve) or *>* 1 (orange curve) which corresponding to a complete sphere or a sphere connected with a spherical cap (Figs. 2E and 2F). This abrupt jump in *η* captures a snap-through transition (*75–77*), which is associated with an abrupt membrane shape transition from a spherical cap to a full sphere bud.

Finally, to understand the interaction between glycopolymer grafting density and length, we constructed two *N*_*p*_− *N* phase diagrams (Fig. 2G corresponding to *κ* = 10 *k*_*B*_*T, λ* = 0 *k*_*B*_*T/*nm and *c*_0_ = 0 nm^−1^ and Fig. 2H corresponding to *κ* = 25 *k*_*B*_*T, λ* = 0.5 *k*_*B*_*T/*nm and *c*_0_ = 0.04 nm^−1^). These phase diagrams show the predicted optimal shape parameter as a function of the number of polymer chains (correlated to grafting density and grafting area) and glycopolymer length to characterize the combined effects of these physical properties of glycocalyx on membrane shape. We observe three regions in the phase diagram corresponding to shallow spherical cap (I: *η*≤ 0.5), deep spherical cap (II: *η >* 0.5), and full sphere (III: *η* = 1). We found that with increasing glycopolymer grafting density or length, the membrane shape shifts smoothly (with a continuous transition) from shallow spherical cap to deep spherical cap when *N*_*p*_ and *N* reach some certain values (blue curves in Figs. 2G and 2H). However, this transition becomes discontinuous from a spherical cap to a full sphere when *N*_*p*_ and *N* exceed certain thresholds shown in the orange dashed curves in Figs. 2G and 2H. These thresholds correspond to the first order transition value that triggers the membrane shape change from a spherical cap to a full spherical bud. Moreover, the height of the glycopolymer brush, *L*_brush_, and the maximum height, *Z*_max_, are also affected by the glycopolymer grafting density and monomer number. Their dependence on glycopolymer grafting density and monomer number are plotted in SI 1 Fig. S2. Thus, we predict that the glycocalyx by itself can generate membrane curvature to minimize the energy of the membrane-polymer composite. Such curvature depends on the density of the glycocalyx and its thickness. These predictions are consistent with previous experimental observations by Shurer et al. (*30*) and simulations by Kutty Kandy et al. (*47*), which indicate that the membrane shape transition requires a threshold polymer density.

### Higher bending rigidity and high membrane tension impede membrane bending

As observed in Figure 2, the shape of the membrane depends on the glycocalyx properties and the membrane properties. To fully understand these dependencies, we investigated the effect of two important membrane parameters: membrane bending rigidity and membrane tension. We first perform the analysis of total free energies for different values of membrane tension and bending rigidity, as shown in Fig. 3A and 3C. Upon decreasing *σ* and *κ*, the energy profiles in Fig. 3A and 3C show similar trends as presented in Fig. 2A-D, indicating that a membrane with lower membrane tension or bending rigidity can trigger a state transition from a spherical cap to a full sphere bud in the case of glycocalyx grafted on a membrane surface. Analogous to Fig 2E, Fig. 3B and 3D show the energetically preferred membrane shape *η*_min_ as a function of membrane tension and bending rigidity. The regime where the optimal shape parameter equals one demonstrates that the globally stable shape is a sphere at low membrane tension and bending rigidity. When tension is increased, the energetically preferred membrane shape *η*_min_ exhibits a sharp transition from *η*_min_ = 1 to *η*_min_ *>* 1, and then slowly increases. While further increases of membrane tension to a critical value will lead to another sharp jump from *η*_min_ *>* 1 to *η*_min_ *<* 1. In the case of *c*_0_ = 0.04 nm^−1^, Fig. 3D displays that as the bending rigidity increases, the energetically preferred membrane shape *η*_min_ exhibits a sharp transition from *η*_min_ = 1 to *η*_min_ *<* 1, and then followed by a slowly decrease. The dependence of brush height and maximum height on membrane bending rigidity and membrane tension are given in SI 1 Fig. S2.

**Figure 3:**
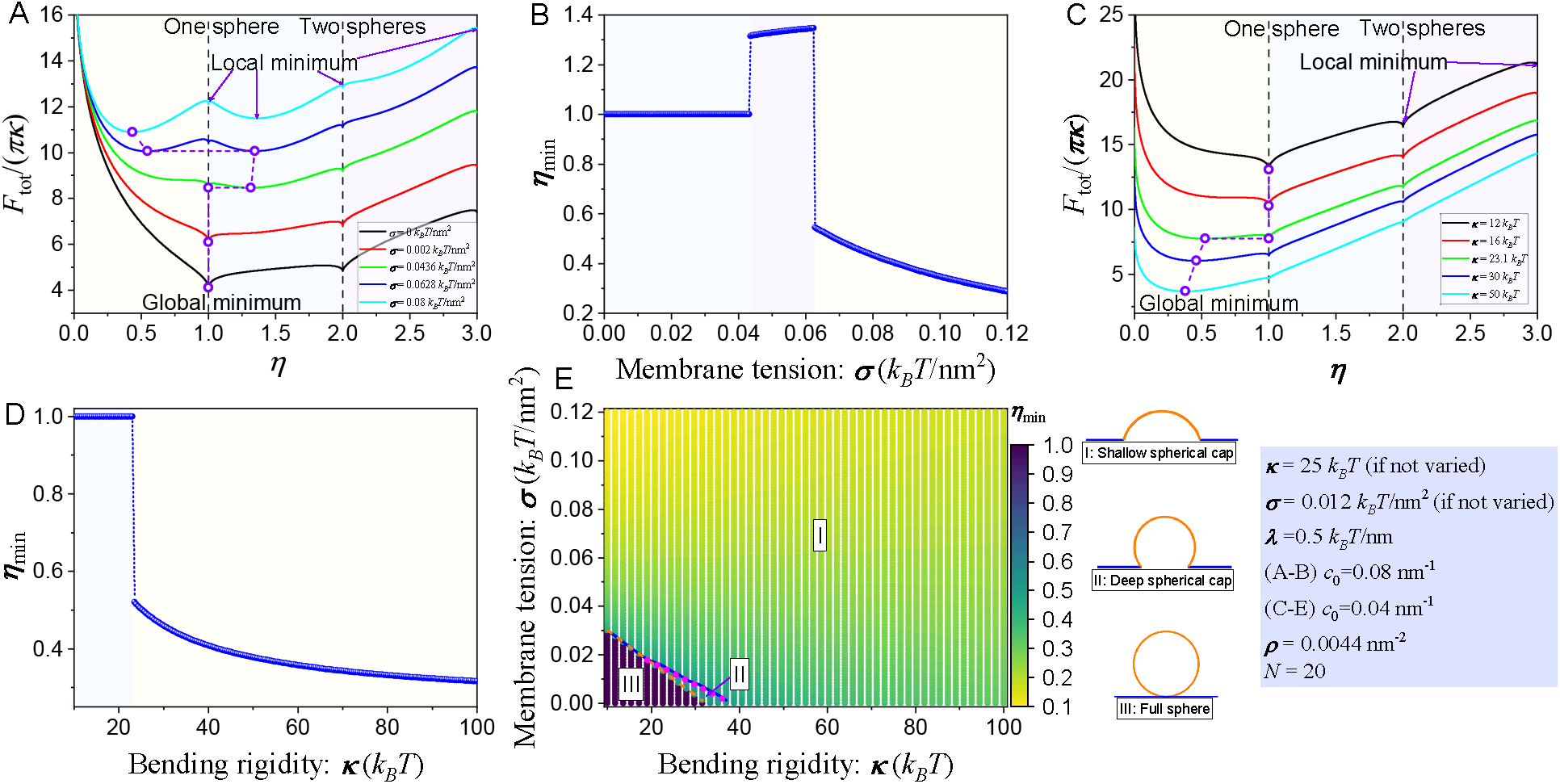
Effects of membrane bending rigidity and membrane tension on membrane reshaping. (A) Total energy profile as a function of shape parameter *η* for different membrane tension *σ*. (B) The dependence of minimum shape parameter *η*_min_ on membrane tension *σ*. (C) Total energy profile as a function of shape parameter *η* for different bending rigidity *κ*. (D) The dependence of minimum shape parameter *η*_min_ on bending rigidity *κ*. (E) A two-dimensional phase diagram on the (*κ*− *σ*) plane characterizes the interrelated effects of bending rigidity and membrane tension on the membrane shape.

The interrelated effects of membrane tension and bending rigidity are summarized in a heatmap (Fig. 3E), where three regions including shallow spherical cap, deep spherical cap, and full sphere are illustrated. Specifically, depending on the membrane rigidity and membrane tension, the cell membrane shape can transition from a shallow spherical cap to a deep spherical cap and eventually to a sphere where the transition from spherical cap to sphere is discontinuous. Our results indicate that higher bending rigidity and a tense membrane impede cell membrane to form a spherical bud. This prediction is in line with the simulation results in Ref. (*47*), where it reported that looser membranes had higher average curvature and that shape transforms in cell membranes in a tension-dependent manner.

### Experiments reveal glycocalyx length-dependent curvature generation

Simulations from our model suggest that steric interactions between glycans within the glycocalyx are sufficient to drive changes in membrane curvature and that the propensity for membrane curvature generation should increase with increasing glycan length and grafting density. To test this hypothesis, we quantified the extent to which overexpression of membrane-bound glycans can induce an increase in the number of positively curved filopodial protrusions at the periphery of adherent mammalian cells. Filopodia are finger-like, actin filled protrusions found at the periphery of cells. They are highly curved, having diameters ranging from 40 to 600 nm (*78*). While actin polymerization is thought to be the main protrusive force responsible for assembly of filopodia (*79*), transmembrane and membrane-bound proteins have also been suggested to contribute (*30*). Here we set out to determine the impact of glycan expression on the assembly of these structures.

For this purpose, we compared the number of filopodia formed per length along the adherent edges of cells (Figure 4), when three model transmembrane proteins were expressed. The model proteins consisted of a non-glycosylated control protein (0TR), a protein which displayed 10 tandem repeats of the Mucin 1, (10TR), and a protein which expressed all 42 native tandem repeats of Mucin 1 (42TR). Each protein was anchored to the plasma membrane by a transmembrane domain and displayed a red fluorescing protein (RFP) domain on the outer surface of the plasma membrane, followed by the prescribed number of tandem repeats (0, 10, or 42). Here the tandem repeat sequence of Mucin 1 is 20 amino acids long and contains 3 threonine residues and two serine residues, each of which are targets of O-glycosylation (refs). Our previous work has established that these model proteins are heavily glycosylated when expressed on the plasma membrane surface, where the degree of glycosylation increases with increasing tandem repeat number (*33*). Fig. 4A shows representative images of cells expressing each of the model proteins. Fig. 4B, shows the average number of filopodial protrusions per length along the perimeter of adherent cells, where the horizontal axis is the expression level of each protein. Notably for cells expressing 10TR and 42TR, the density of filopodia increased with increasing expression of the proteins (Fig. 4B). Fig. 4C summarizes these data, demonstrating that the expression of 10TR and 42TR progressively increased the density of filopodia relative to cells expressing 0TR. Our experimental results show that the filopodia number exhibits an increasing trend with respect to native tandem repeats, as shown in Fig. 4, indicating that thicker glycocalyx generates more curvature on the membrane.

**Figure 4:**
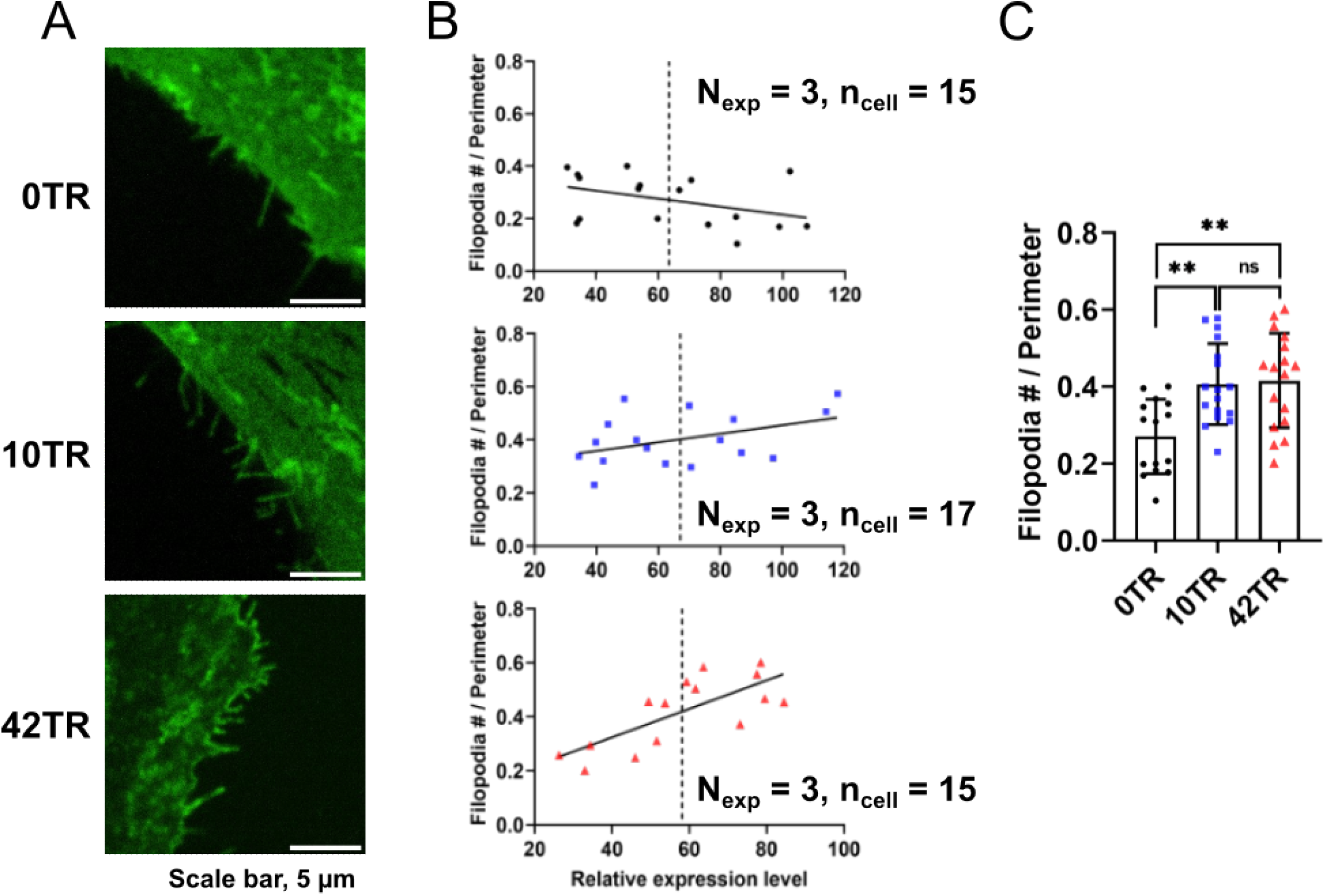
Experimental results of curvature generation capabilities. Glycosylated proteins contribute to the increasing number of filopodia. (A) Representative fluorescent images and (B) corresponding plots showing the relationship between the number of Filopodia to cell perimeter versus the relative expression level of GFP. *N*_exp_ represents an independent experimental repetition and *n*_cell_ represents the number of cells to be measured. (C) Bar plots of average number of filopodia. ns indicates no significant difference; ^∗∗^*P <* 0.01 by one-way ANOVA with Tukey’s test.

### Interplay between spontaneous curvature, line tension, and the glycocalyx

We next wondered under what conditions might we observe a richer diversity of membrane shapes from our model. For example, many experimental studies have reported that the glycocalyx (*30*) and other polymers (*44, 80, 81*) can generate pearled morphologies or beads-on-a-string structures. To understand the physics behind these shapes, we altered the spontaneous curvature and line tension (Fig. 5 and Fig. S3). In the absence of line tension, we find that the shape corresponding to an energy minima shifts from 0 *< η*_min_ *<* 1 towards *η*_min_ *>* 1 with increasing spontaneous curvature (Fig. 5A). The same trend is found for the energy profile when the line tension is non-zero, as shown in Fig. 5D. However, the difference between them is that *η*_min_ can be non-integer at zero line tension, while *η*_min_ is only an integer when the line tension is greater than a certain value. To evaluate this difference, we analyze the energy variations of different contributions (Fig. 5B, E). The reason is attributed to the energy kinks arising from the non-smooth transitions of membrane tension energy and line tension energy at integer *η*. At these points, The interface of glycocalyx-rich and glycocalyx-poor phase will disappear and the base radius, *R*_*b*_, becomes zero when a full sphere is formed at integer *η*. When line tension is zero, the energy profile of membrane tension energy exhibits multiple local minima with corresponding shape parameters that are non-integer (Fig. S4B). As a result, this effect gives rise to a non-integer *η*_min_ of the global local minimum. However, when the line tension is included, the local minima of line tension energy are located where the shape parameters are integer (Fig. 5E and Fig. S4C). This consequently yields an integer *η*_min_ of the global or local minimum. The scenario of how *η*_min_ depends on spontaneous curvature is illustrated in Fig. 5C and Fig. 5F, respectively. As the spontaneous curvature is increased, one obtains stable pearled shapes with a series of equal spheres connected to a spherical cap at zero line tension (Fig. 5C), and a stack of equal spheres structure in the presence of line tension (Fig. 5F). Also, the energetically preferred membrane shapes undergo a discontinuous transition. The dependence of brush height and maximum height on spontaneous curvature are displayed in SI 1 Fig. S2.

**Figure 5:**
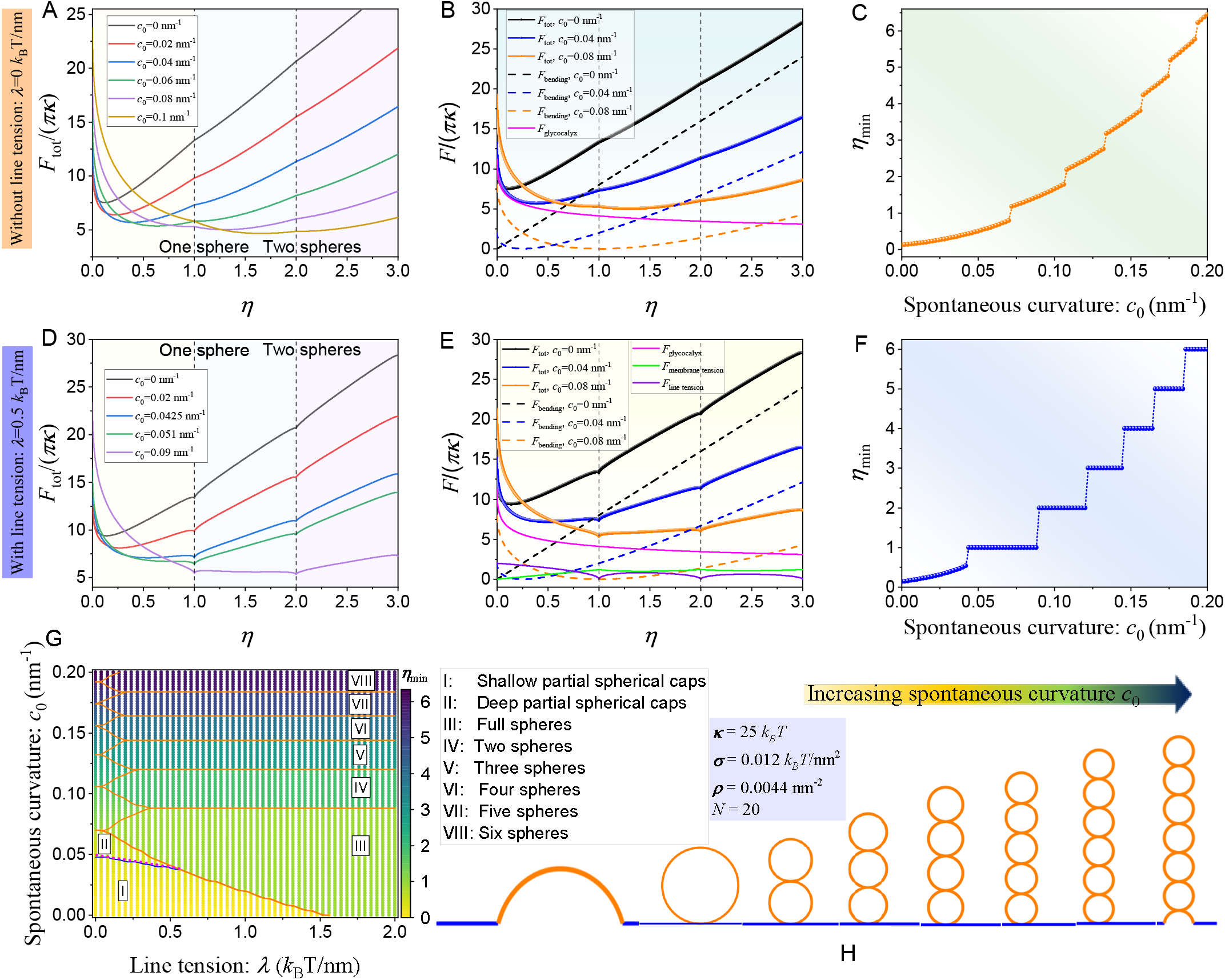
Sensitivity of membrane curvature to spontaneous curvature and line tension. (A) Total energy and (B) component energy contributions profiles as a function of shape parameter *η* for different spontaneous curvature *c*_0_ in the absence of line tension. (C) The dependence of minimum shape parameter *η*_min_ on spontaneous curvature *c*_0_. (D) Total energy and (E) component energy contributions profiles as a function of shape parameter *η* for different spontaneous curvature *c*_0_ when the line tension is fixed at *λ* = 0.5 *k*_*B*_*T/*nm. (F) The dependence of minimum shape parameter *η*_min_ on spontaneous curvature *c*_0_. (G) A two-dimensional phase diagram on the (*λ*− *c*_0_) plane characterizes the interrelated effects of line tension and spontaneous curvature on the membrane shape. (H) Various membrane morphologies correspond to spherical cap, sphere, and pearled shapes.

These findings are summarized in a two dimensional phase diagram (Fig. 5G), where a variety of regions including shallow spherical cap, deep spherical cap, full sphere and different types of pearled structure are illustrated. Specifically, depending on the spontaneous curvature and line tension, the cell membrane shape can transit from a shallow spherical cap to a deep spherical cap and eventually to a sphere where the transition from spherical cap to sphere is discontinuous. Interestingly, the occurrence of a pearled membrane structure was found to largely depend on the spontaneous curvature of the membrane. Figure 5H shows several typical cell membrane morphologies by increasing the spontaneous curvature. The energy profile in Fig. S3A depicts that the energetically more favorable state corresponding to membrane shape 0 *< η*_min_ ≤ 1 at small spontaneous curvature and *η*_min_ ≥ 1 at large spontaneous curvature as line tension is increased. Meanwhile, the dependence of minimum shape parameter *η*_min_ on line tension *λ* are displayed in Fig. S3, the dependence of brush height and maximum height on line tension *λ* are provided in Fig. S2. As predicted by Eq. S16, the brush height and the maximum height decreases and increases, respectively, with the increasing of the number of equal sphere structure generated via the increasing of spontaneous curvature. Thus, our model predicts that membrane spontaneous curvature plays a significant role in the formation of pearled membrane structures.

### Heterogeneity of glycocalyx distribution as a function of membrane curvature

Thus far, we have only considered curvature generation induced by the glycocalyx that is confined to a specific region of the membrane. However, the dependence of the spatial heterogeneity distribution of glycocalyx on local membrane curvature is still not fully understood. To explore this curvature dependence of glycocalyx, we developed a thermodynamically consistent model for heterogeneity of glycocalyx distribution. We also carried out experiments to study the relative distribution of glycocalyx on cell membrane surface, and further to test our theoretical predictions. To quantify the curvature sensing ability of glycocalyx, based on the thermodynamic model developed above, we define a relative partitioning number as *ϕ*_*t*_*/ϕ*_*V*_, where *ϕ*_*t*_ is the area fraction of glycocalyx on the tube, and *ϕ*_*V*_ is the area fraction on the cell surface, as shown in Fig. 6A. Based on our model, we calculated the effects of glycopolymer length on the relative partitioning number. As shown in Fig. 6B, our theoretical model predicts an increase in relative partitioning with increasing polymer length. This trend reflects the curvature sensing ability of the glycocalyx. Note that the glycocalyx prefers higher local membrane curvature and this curvature dependence refers to curvature sensing. Thus, our model predicts that the glycocalyx partitions to higher curvature indicating its curvature sensing capability.

**Figure 6:**
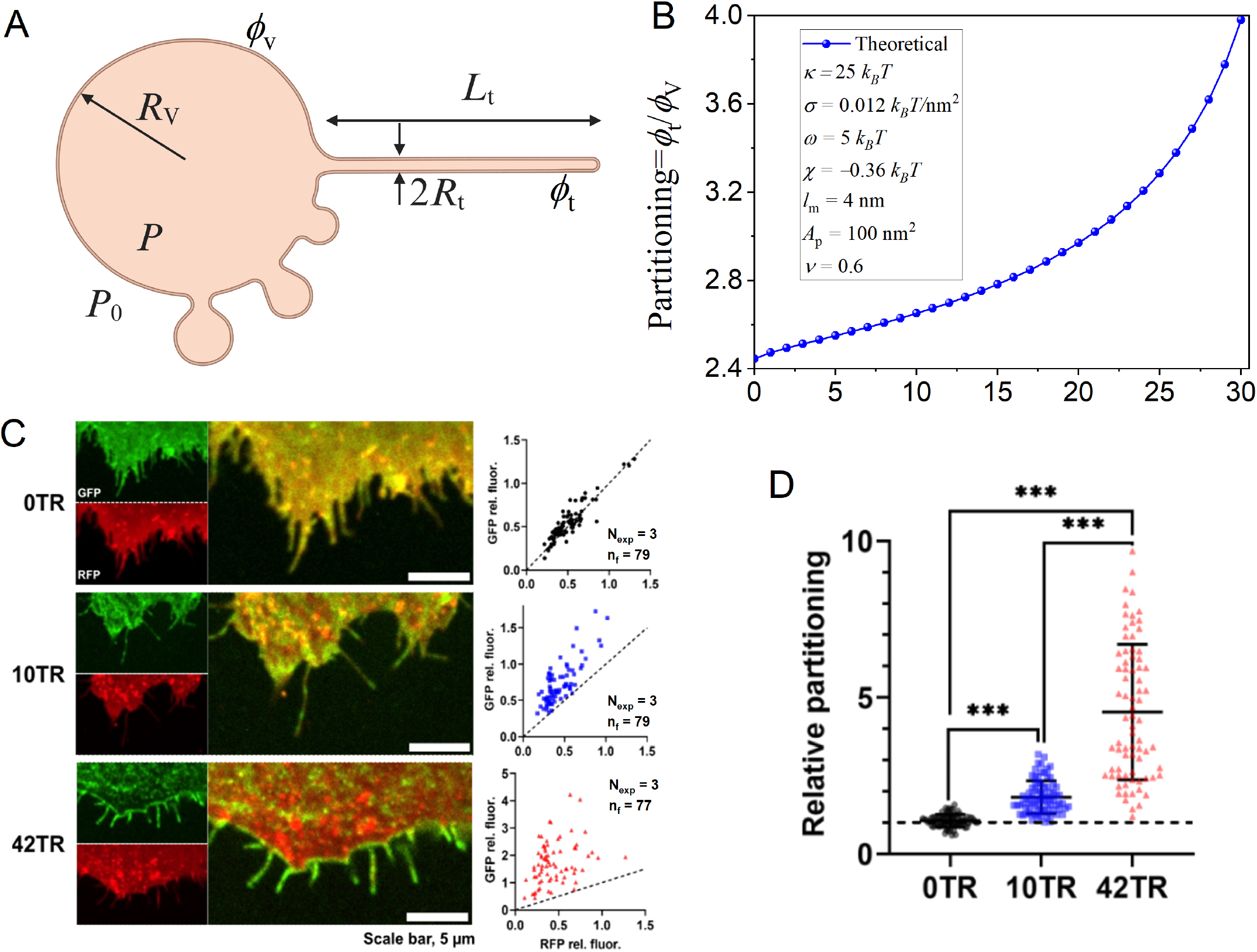
Curvature sensing of glycocalyx on cell membrane. (A) Schematic of the system as used in the curvature sensing model. We consider the membrane to consist of a vesicle component that represents the whole cell and tubular component with radius *R*_*t*_ and length *L*_*t*_. (B) Theoretical prediction of the ratio between the area fraction of polymer on the tube and on the cell surface as a function of polymer length. Glycosylated proteins sense the curvature of filopodia in live cells. (C) Fluorescence microscopy images of filopodia, along with plots of relative GFP versus RFP fluorescence on filopodia for 0TR, 10TR, and 42TR, respectively. *N*_exp_ represents independent experimental repetitions, and *n*_f_ represents the number of filopodia to be measured. The dashed line corresponds to no curvature sensitivity. (D) Bar plots showing the average change in filopodia partitioning for GFP-labeled proteins relative to RFP-labeled control proteins. ^∗∗∗^*P <* 0.01 by one-way ANOVA with Tukey’s test.

To test these predictions, we returned to the experimental system shown in Fig. 4 above, which used model transmembrane proteins with different numbers of O-glycosylatable mucin 1 tandem repeats: 0TR, 10TR, 42TR. To assess the ability of these proteins to sense membrane curvature, we measured their partition coefficient between positively curved filopodia and the plasma membrane, which is relatively flat on average. Specifically, we co-expressed the GFP-labeled model proteins (0TR, 10TR, and 42TR) alongside an RFP labeled control protein that lacked tandem repeats, RFP-0TR. Co-expression of this control protein, which would not be expected to sense membrane curvature, provided a reference point from which to judge curvature sensing by the GFP-labled model proteins. We measured the intensity in the GFP and RFP channels along each filopodium and on the plasma membrane near the base of each filopodium. To assess curvature sensing, we plotted the ratio of GFP intensity to RFP intensity for the filopodium on the vertical axis and the ratio of GFP intensity to RFP intensity on the plasma membrane on the horizontal axis, as shown in Fig. 6C. If the data on this plot were to fall along a line of slope 1, it would indicate that the GFP-labeled receptor (0TR, 10TR, or 42TR) partitioned between filopodia and the plasma membrane equally with the partitioning of RFP-0TR. As RFP-0TR is presumed to lack sensitivity to membrane curvature, such a result would indicate a lack of curvature sensing ability by the GFP-labeled protein. Examining Fig. 6C, it is clear the data for cells expressing the GFP-labeled 0TR protein has a slope very close to one, indicating a lack of curvature sensitivity, as we would expect in the absence of any glycosylatable tandem repeats. In contrast, the corresponding plots for cells expressing the 10TR and 42TR model proteins fall well above the line of slope equal to one, indicating that these protein partition substantially more favorably to filopodia in comparison to the RFP-0TR control protein. Figure 6D shows the average partitioning values for each of the three proteins, 0TR, 10TR, and 42TR. These data indicate a substantial increase in partitioning to filopodia with increasing tandem repeat length. Notably, the 42TR model protein, which displays the full Mucin1 tandem repeat domain, favors filopodia over the plasma membrane by a factor of about five to one.

## Discussion

In this study, we have established a mechanical framework to explain how the Helfrich energy and the energy of the glycocalyx can interact to regulate membrane curvature. We show that the glycocalyx is capable of generating membrane curvature even in the absence of an intrinsic spontaneous curvature indicating that the glycocalyx alone is able to generate outward membrane bending. Furthermore, we show that increasing of glycopolymer density and length can yield membrane buds similar to those observed experimentally (*30*) (see Fig. 2). Having identified the glycocalyx properties as key control variables, we tested the role of number of mucin TRs in cells and showed that the number of filopodia on cells depend on the length of the glycocalyx (Fig. 4). Finally, we used theory and experiments to quantify the curvature sensing ability of glycocalyx. We predicted that the increasing of polymer length leads to a increase in partitioning number. This prediction was verified experimentally and indicates that the glycocalyx prefers higher local membrane curvature (see Fig. 6). Additional analyses show that a spherical shape is favored for lower membrane bending rigidity and membrane tension even in the presence of the glycocalyx (see Fig. 3) and that the spontaneous curvature is a key control parameter for the generation of pearl-shaped membrane structure (see Fig. 5). Thus, we conclude that the membrane shape is determined by the competition among the elastic energy (consisting of bending energy and tension), the energy contribution associated with the glycocalyx, and line tension energy.

Our findings have implications for different membrane trafficking processes. Outward bending of the membrane (away from the cytosol) is encountered in filopodia formation, blebbing, and mircroparticle formation (*79, 82–84*). While the role of the cytoskeleton and membrane lipids have been investigated in these studies (*84–88*), the specific role of the glycocalyx has not be elucidated in these processes. It is possible that the glycocalyx, in fact, inherently provides an energetically favorable state to bend outward. It is also possible that the favorable energy contribution of the glycocalyx to outward bending can overcome the actin-membrane cortex attachment in cases where blebs or extracellular vesicles need to form. This idea is also consistent with studies showing that the glycocalyx opposing inward budding as in the case of endocytosis (*33*). Indeed, the direction of inward or outward budding can be controlled by the net stresses on the membrane (*34*). Here, we show that the glycocalyx has specific ways, including length and density control, through which these stresses can be controlled to promote outward bending.

Recent studies have shown that the structure and organization of the glycocalyx are altered in most cancer cells, which can result in modified binding characteristics and downstream signaling (*36, 89*). Changes in the bulk physical properties of glycocalyx can affect the cellular morphologies through multiple mechanisms that cause cell membrane remodeling, which can alter the formation of extracellular vesicles and exosomes that regulate cell-cell communication. Enzymes that promote glycocalyx degradation often result in the loss of the thickness of the glycocalyx layer and its mechanical integrity (*90–93*). Inflammation-mediated shedding of different glycocalyx molecules can lead to disrupted endothelial signaling, vascular hyperpermeability, and unregulated vasodilation among other conditions (*91, 92*). Our modeling effort opens up the possibility of investigating the role of changes to the glycocalyx thickness in cell function under physiological and pathological conditions.

We note that future modeling and experimental studies need to incorporate additional biological details of the cellular plasma membrane beyond the lipid bilayer to dissect the contributions of different moieties on the curvature directions. Our current calculations in our work relied on certain restricted geometries but future efforts can extend these theories to new computational frameworks (*94, 95*). Future efforts also need to consider the spatial heterogeneity of the distribution of the glycocalyx and the effects of charged polymers to capture the biological complexity of cell membranes.

## 2 Acknowledgments

This work was supported by NIH R01GM132106 to P.R, NSF MCB 2327243 to P.R. and J.C.S., and NIH R35GM139531 to J.C.S. We would like to thank Dr. Aravind Chandrasekaran and Mr. Nathaniel Linden for their thoughtful feedback. The authors have no conflicts of interest to declare.

## 3 Author contributions

K.X., J.C.S, and P.R. conceived the work. K.X. and P.R. developed the analytical model, X.K. conducted the simulations, S.P. conducted the experiments. J.C.S and P.R. are responsible for overseeing the work and obtaining funding. All authors contributed to the writing and revision of the manuscript.

## Supporting Information (SI)

We provide here a detailed description of the mathematical methods we used for the calculation of the energy contribution of glycocalyx polymers and associated membrane deformation energy.

### 1 Curvature Generation Model: Spherical cap geometry

As explained in the main text, we calculated the membrane of glycocalyx attached by finding the stationary membrane shapes associated with the classic Helfrich energy (*48*)

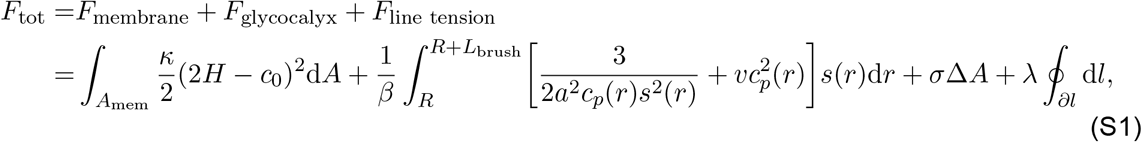

where *β* = 1*/*(*k*_*B*_*T*) with *k*_*B*_*T* being the thermal energy.

#### 1.1 Description of spherical cap shape

To estimate the energy described in Eq. (S1), we assume that the membrane patch grafted with glycocalyx polymer brushes takes the shape of spherical caps with radius *R*_*s*_, opening angle 2*α*, and fixed surface area 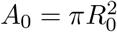, where *R*_0_ is the in-plane radius of the flattened membrane patch. Based on experimental observations (*30*), during the membrane deformation process, we assume the patch of the membrane with glycocalyx polymers on it is gradually bent (assisted by glycocalyx polymers) into the shape of spherical caps (Fig. 1C in the main text). This restricted family of shapes are conveniently specified by a shape parameter *η* which is defined as the fraction of the surface of spheres of uniform radius *R*_*s*_ covered by the glycocalyx polymers, 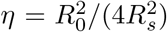,with *η* = 0 corresponding to a completely flattened membrane, 0 *< η <* 1 for the membrane taking the shape of shallow or deep spherical caps, *η* = 1 for a full sphere, and *η >* 1 corresponding to pearled intermediate with *n* = [*η*] spheres. These special geometries also allow us to express the radius and opening angle 2*α* of the spherical cap as a function of the shape factor *η*, which are given by 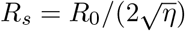 and cos *α* = 1 − 2*η* + 2*n*, respectively.

#### 1.2 Elastic energy of cell membrane

Specifically, we treat the cell membrane as a homogeneous thin elastic layer. To describe the energy cost of bending the membrane away from its preferred shape, we use the well-known Canham-Helfrich energy density (*48*), *f*_*b*_ = *κ*(2*H*− *c*_0_)^2^*/*2, where *κ* is the membrane bending rigidity, *H* = (*c*_1_ + *c*_2_)*/*2 is the mean curvature in which *c*_1_ and *c*_2_ are the two principal curvatures, and *c*_0_ is the spontaneous curvature, respectively. Here, in this model, *c*_1_ = *c*_2_ = 1*/R*_*s*_ because of the spherical geometry. Thus, the energy contribution due to membrane bending, *F*_*b*_, can be then obtained as an integral of *f*_*b*_ over the entire surface, which is calculated as

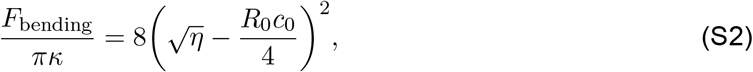

where dividing by *πκ* normalizes the energy.

Remodeling the membrane away from an initially flat shape also involves a surface energy contribution, *F*_tension_ = *σ*Δ*A*, which corresponds to the work required to form a curved membrane shape against a lateral membrane tension *σ*, where Δ*A* denotes the excess area compared to a flat membrane. For the spherical cap and a series of unduloids geometries, the excess area is defined as 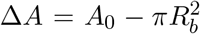 with the in-plane base radius 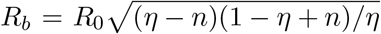. Therefore the contribution of the tension to the energy is

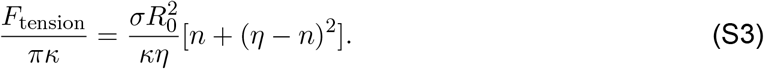

#### 1.3 Line tension energy

We also consider the interfacial energy between a glycocalyx rich domain and a bare lipid layer, *F*_line tension_ = *λ* ∮_*∂l*_ d*l*, where *λ* represents the strength of line tension along the domain boundary, and the integral is over the periphery line d*l* of the domain. Such a nonzero line tension can arise, for example, from differences in molecular composition between transmembrane domains of the glycocalyx and the surrounding membrane (*96*). Interactions between glycocalyx polymers and the surrounding membrane may also yield an effective contribution to the line tension that depends on membrane morphology. We assume that the line tension is a constant. Based on the prescribed membrane geometry, the line tension energy is given by

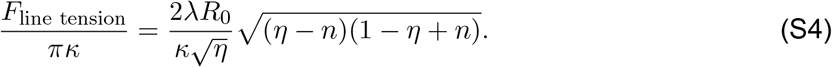

#### 1.4 Energy contribution of glycocalyx polymer

Notably, the crowding of large glycosylated proteins appears to regulate the shape of the underlying bilayer plasma membrane (*30*). According to polymer physics, two regimes can be defined depending on the grafting density of glycocalyx polymers on the cell membrane, which are the mushroom-like regime and the brush-like regime. Shurer et al. (*30*) have reported that the mucins are in the coiled mushroom state in the case of low-density glycocalyx polymers. We focus on densely grafted regions of the membrane and therefore model the glycocalyx in the brush regime.

Here, we consider a layer of glycocalyx grafted on a membrane of radius *R*_*s*_ in a brush-like structure with thickness *L*_brush_, as shown in Fig. S1. From the viewpoint of coarse graining, the polymer brush is envisioned as an array of blobs. The size of each blob, *ξ*, at a given position *r*, equals the square root of the local area per chain *s*(*r*), where *r* = *x* + *R*_*s*_ is the radial distance which is defined from the center of the spherical surface, and in which *x* is the distance from the membrane surface. Thus, the blob size 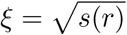 grows as a function of *r*, and the grafting density of glycocalyx polymer brush on the membrane surface can be obtained as *ρ* = 1*/ξ*^2^. Assuming that the layer of glycopolymers is extended non-uniformly but equally in height, *L*_brush_, then the area per chain at distance *x* from the membrane surface is given by (*49*)

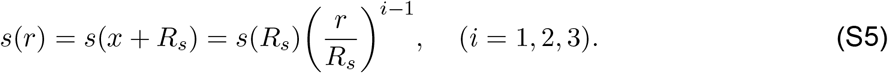

The index *i* = 1, 2, 3 indicates planar, cylindrical, and spherical shaped membranes, respectively. When the membrane is bent, the changes of polymer configuration give rise to the local extension of the polymer chain. Here the local chain extension at a height *r* is characterized by d*r/*d*n*_*b*_, where the variable *n*_*b*_ denotes the current monomer. This local extension is related to the local density profile of monomers *c*_*p*_(*r*) as (*49*)

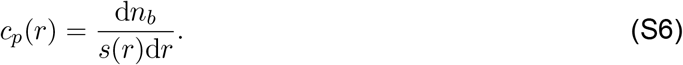

**Figure S1:**
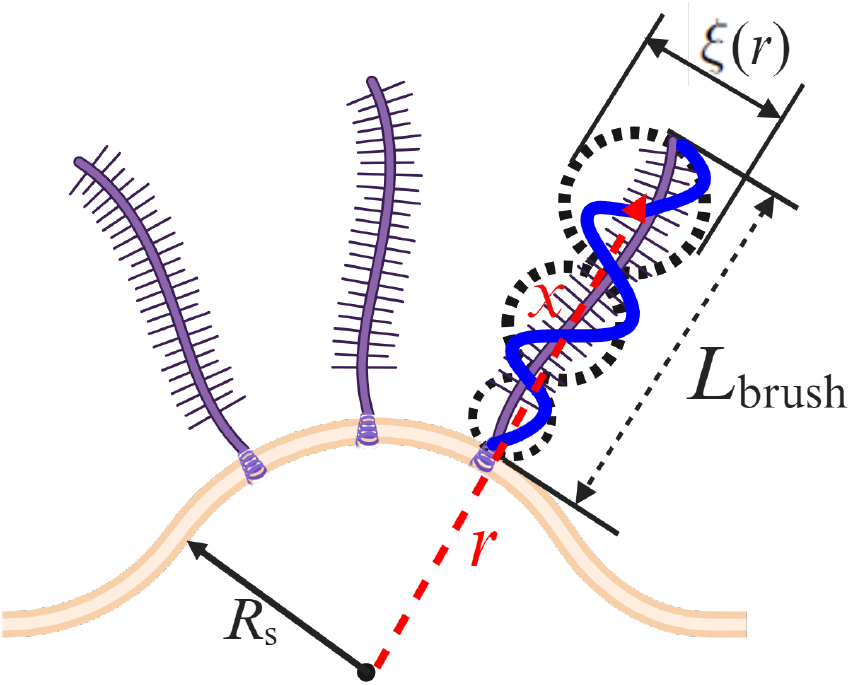
Schematic of a layer of brush-like polymer attached on a spherical cap membrane surface with radius *R*_*s*_. At a given position *r*, the coarse grained blob (dashed black circle) size in a polymer brush is *ξ*(*r*), and the thickness of the brush is *L*_brush_.

Then, the thickness of the brush, *L*_brush_, is found from the conservation condition (the constraint of conservation of the total number of monomers *N*) (*49*),

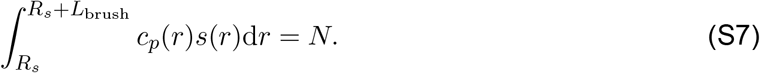

As a result, in the brush regime, since the electrostatic effects are excluded, the energy contribution original from the glycocalyx polymers includes two terms (*30, 49*): the elastic energy of the polymer chain 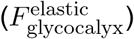 and the free energy caused by the excluded volume interactions of polymer monomers 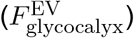. Based on the hypotheses in the main text, according to Ref. (*49*), the elastic energy per chain in the brush can be represented as

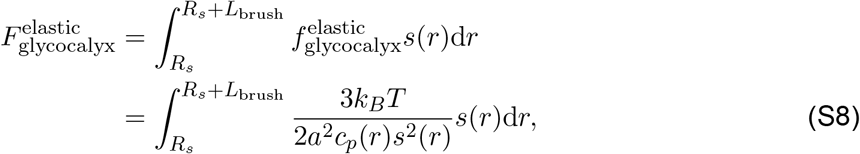

where *a* is the monomer length and 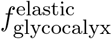 is the elastic energy density of the polymer chain. Within the mean-field approximation, the energy density of the excluded volume interactions (van der Waals interactions) between monomers can be modeled in terms of the virial expansion

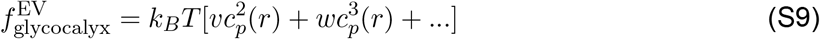

where *v* and *w* are the second and third virial coefficient, respectively. As a result, the excluded volume interactions between monomers per chain in the brush is given by

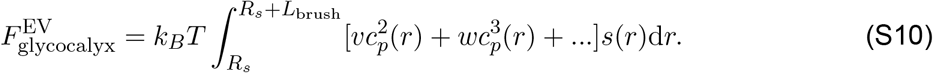

In subsequent analysis, the cubic and higher terms are neglected. Therefore, the sum of Eq. (S8) and Eq. (S10) yields the energy contribution of glycocalyx polymers

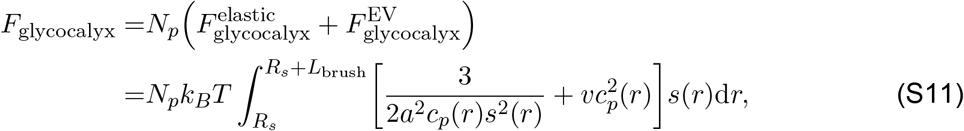

where *N*_*p*_ = *ρA*_0_ is the number of polymer chains grafted on the membrane. Here *A*_0_ stands for the membrane patch area covered by glycocalyx polymers and *ρ* is the density.

In order to calculate the free energy, we need to further determine the local concentration of monomers *c*_*p*_(*r*) and the brush thickness *L*_brush_. On a planar membrane surface, the planar brush area per chain, *s*(*r*), is constant, i.e., *s*(*r*) = *s*. Minimizing the free energy of a planar brush *F*_glycocalyx_ with respect to *c*_*p*_ by using 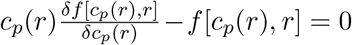 leads to the equilibrium polymer Concentration 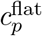 (see Ref. (*49*) for detailed steps)

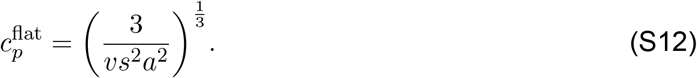

Here, 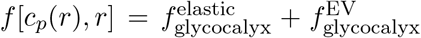. Using the conservation condition defined by Eq. (S7) yields the brush thickness 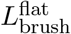 (see Ref. (*49*) for detailed steps)

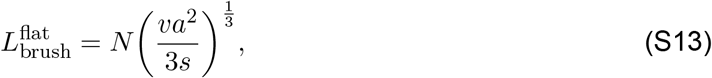

where we can see that the thickness of the polymer brush is proportional to the total number of monomers *N*. Hereafter, in our model, we use the number of monomers, *N*, to capture the length of the polymer. Note that the relationship for 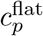 and 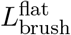 are special cases of electrically neutral brushes as described in (*49*). Substituting Eq. (S12) and Eq. (S13) into Eq. (S11) yields the energy contribution of glycocalyx polymers on a planar membrane surface

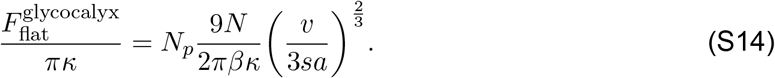

Thus, even for a flat membrane, the energy contribution by the glycocalyx is directly proportional to the extent of grafting *N*_*p*_ and the length of the polymer brush *N*.

On a spherical membrane surface, the corresponding monomer density profile *c*_*p*_(*r*) and brush thickness *L*_brush_ are, respectively, expressed as (see Ref. (*49*) for a detailed calculation)

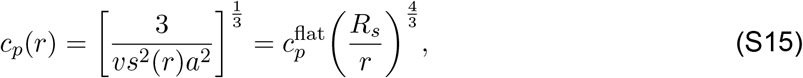

And

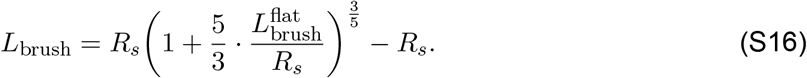

Similarly, substituting Eq. (S15) and Eq. (S16) into Eq. (S11) leads to the energy contribution of glycocalyx polymers on a spherical membrane surface as

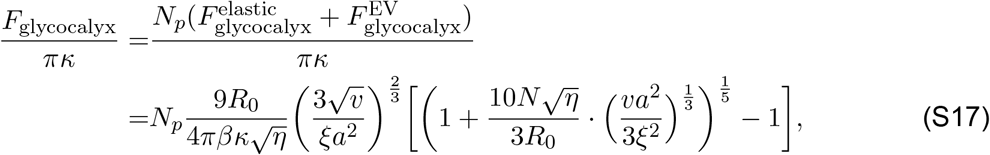

where we use 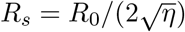.

#### 1.5 Total energy of the system

Combining Eq. (S2), Eq. (S3), Eq. (S4), Eq. (S14), and Eq. (S17) leads to the Eq. (5) in the main text.

To obtain an analytical expression for the boundary curves that separate the shallow spherical cap regime from the deep spherical cap regime, as shown by the dotted magenta curves in the phase diagrams in Figs. 2, 3, and 5 in the main text. Taking a first order derivative of the total energy *F*_tot_ with respect to *η*, one can obtain

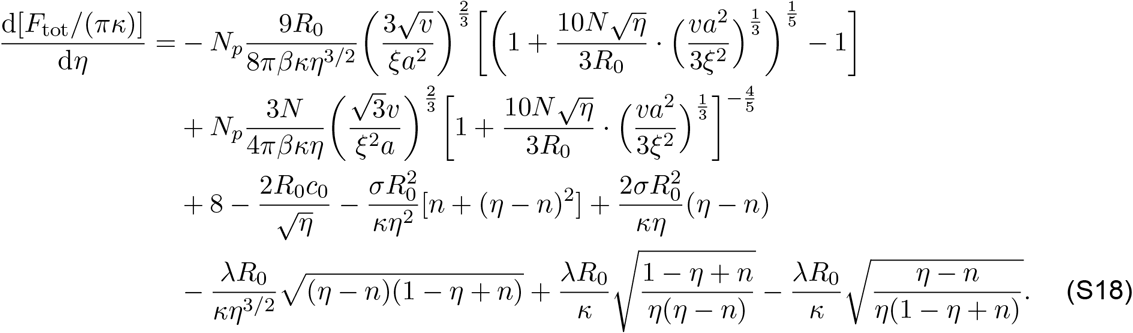

By setting d[*F*_tot_*/*(*πκ*)]*/*d*η* = 0, we can get the analytical solution that corresponds to the critical transition line between the shallow spherical cap regime and the deep spherical cap regime, which is shown by the dotted magenta curves in the phase diagrams in Figs. 2, 3, and 5 in the main text.

The values of the parameters used in each figure in the main text are listed as follows. In Fig. 2A, 2B, 2E(orange line), 2F(orange line)and 2G, the parameters are fixed at: *κ* = 10 *k*_*B*_*T, σ* = 0.012 *k*_*B*_*T/*nm^2^, *λ* = 0 *k*_*B*_*T/*nm, *c*_0_ = 0 nm^−1^, *ξ* = 11.9 nm (*ρ* = 0.007 nm^−2^ if not varied), and *N* = 30 (if not varied). In Fig. 2C, 2D, 2E(blue line), 2F(blue line)and 2H, the parameters are fixed at: *κ* = 25 *k*_*B*_*T, σ* = 0.012 *k*_*B*_*T/*nm^2^, *λ* = 0.5 *k*_*B*_*T/*nm, *c*_0_ = 0.04 nm^−1^, *ξ* = 15 nm (*ρ* = 0.0044 nm^−2^ if not varied), and *N* = 20 (if not varied).

In Fig. 3, the parameters are fixed at: *κ* = 25 *k*_*B*_*T* (if not varied), *σ* = 0.012 *k*_*B*_*T/*nm^2^ (if not varied), *λ* = 0.5 *k*_*B*_*T/*nm, *ξ* = 15 nm (*ρ* = 0.0044 nm^−2^), *N* = 20, and *c*_0_ = 0.08 nm^−1^ for Fig. 3A and 3B, and *c*_0_ = 0.04 nm^−1^ for Fig. 3C-3E.

In Fig. 5, the parameters are fixed at: *κ* = 25 *k*_*B*_*T, σ* = 0.012 *k*_*B*_*T/*nm^2^, *ξ* = 15 nm (*ρ* = 0.0044 nm^−2^), *N* = 20, and *λ* = 0 *k*_*B*_*T/*nm for Fig. 5A-5C, and *λ* = 0.5 *k*_*B*_*T/*nm for Fig. 5D-5F.

Based on the model developed above, we can also study how glycocalyx properties (including grafting density and monomer number), membrane properties (including bending rigidity and membrane tension), spontaneous curvature, and line tension affect the height of glycopolymer brush *L*_brush_ and the maximum height *Z*_max_ of a deformed cell membrane. To intuitively see the effect of glycocalyx properties, it is helpful to plot the height of glycopolymer brush *L*_brush_ and the maximum height *Z*_max_ against grafting density and monomer number, as illustrated in Fig. S2. As grafting density *ρ* or monomer number *N* increases, *L*_brush_, and *Z*_max_ are shifted smoothly and then followed by a discontinuous transition which corresponding to the membrane shape transition from a spherical cap with 0 *< η <* 1 to a sphere with *η* = 1. We can clearly see that the trend is analogous to the dependence of *η*_min_ on glycocalyx properties.

To study the influence of line tension on membrane reshaping, the total energy against shape parameter at different line tension, and the dependence of minimum shape parameter *η*_min_ on line tension are plotted in Fig. S3. To reveal the mechanisms of cell membrane bending regulated by glycocalyx, the calculated total energy and different energy contributions as a function of shape parameter are shown in Fig. S4, demonstrating that membrane bending is governed by a balance among these energy players.

**Figure S2:**
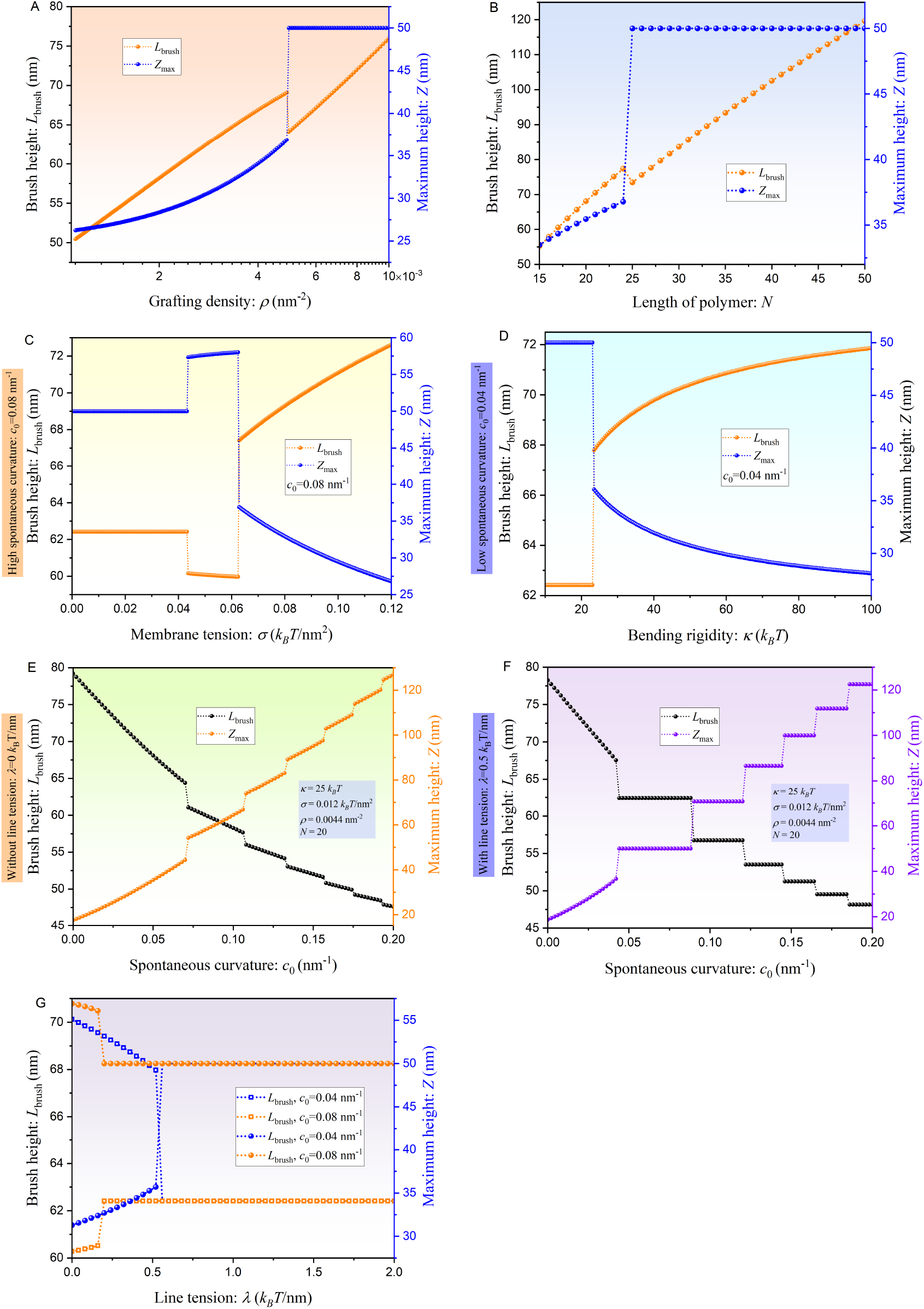
Dependence of membrane height and thickness of brush layer on various parameters. The dependence of brush height *L*_brush_ and maximum height *Z* on (A) grafting density *ρ*, (B) polymer length *N*, (C) membrane tension *σ*, (D) bending rigidity *κ*, spontaneous curvature *c*_0_ (E) without line tension (F) with line tension, and (H) line tension *λ*, respectively.

**Figure S3:**
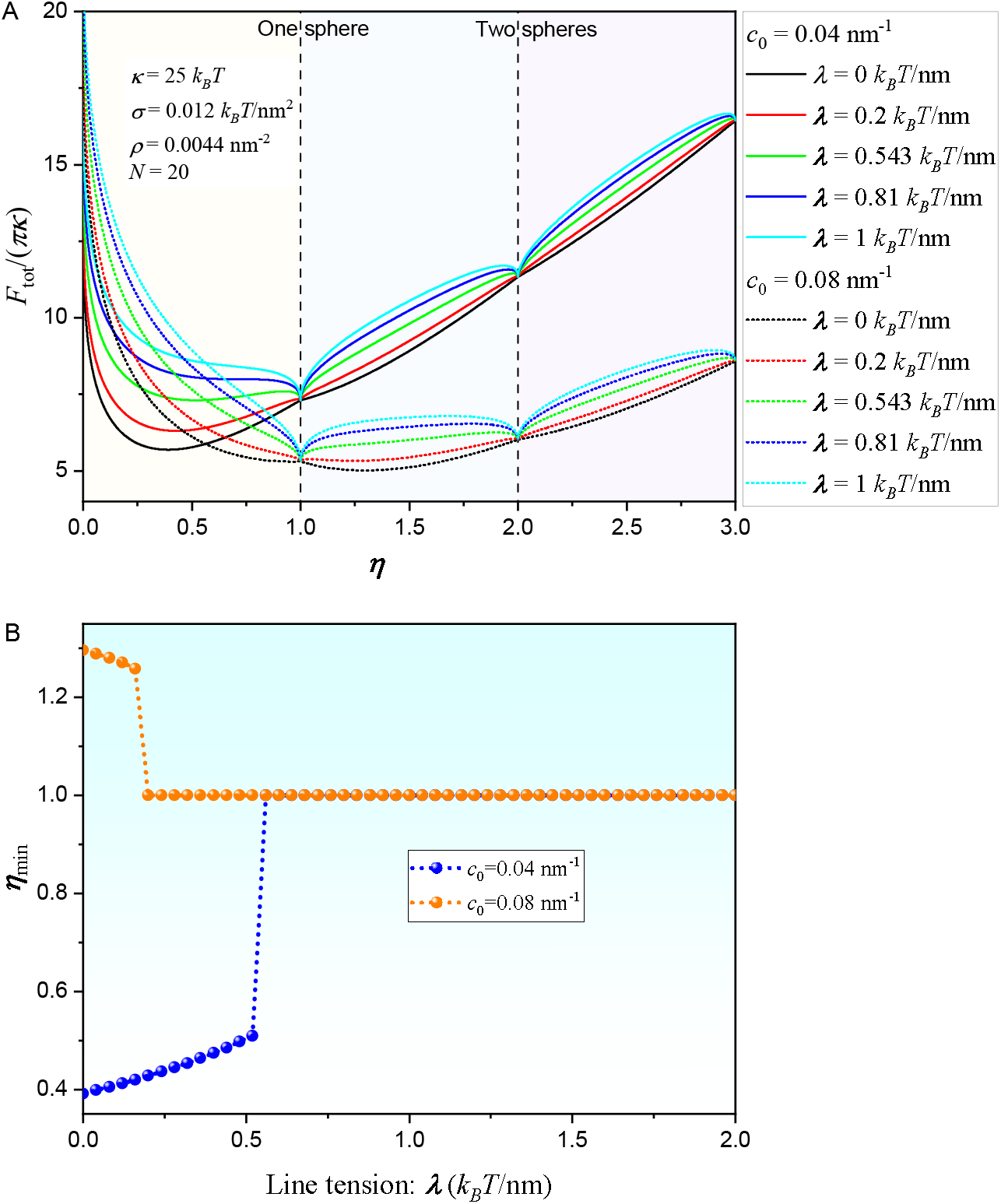
Effects of line tension on membrane shape remodeling. (A) Total energy profile as a function of shape parameter *η* for different line tension *λ* when the spontaneous curvature *c*_0_ are fixed at *c*_0_ = 0.04 nm^−1^ and *c*_0_ = 0.08 nm^−1^, respectively. (B) The dependence of minimum shape parameter *η*_min_ on line tension *λ*.

**Figure S4:**
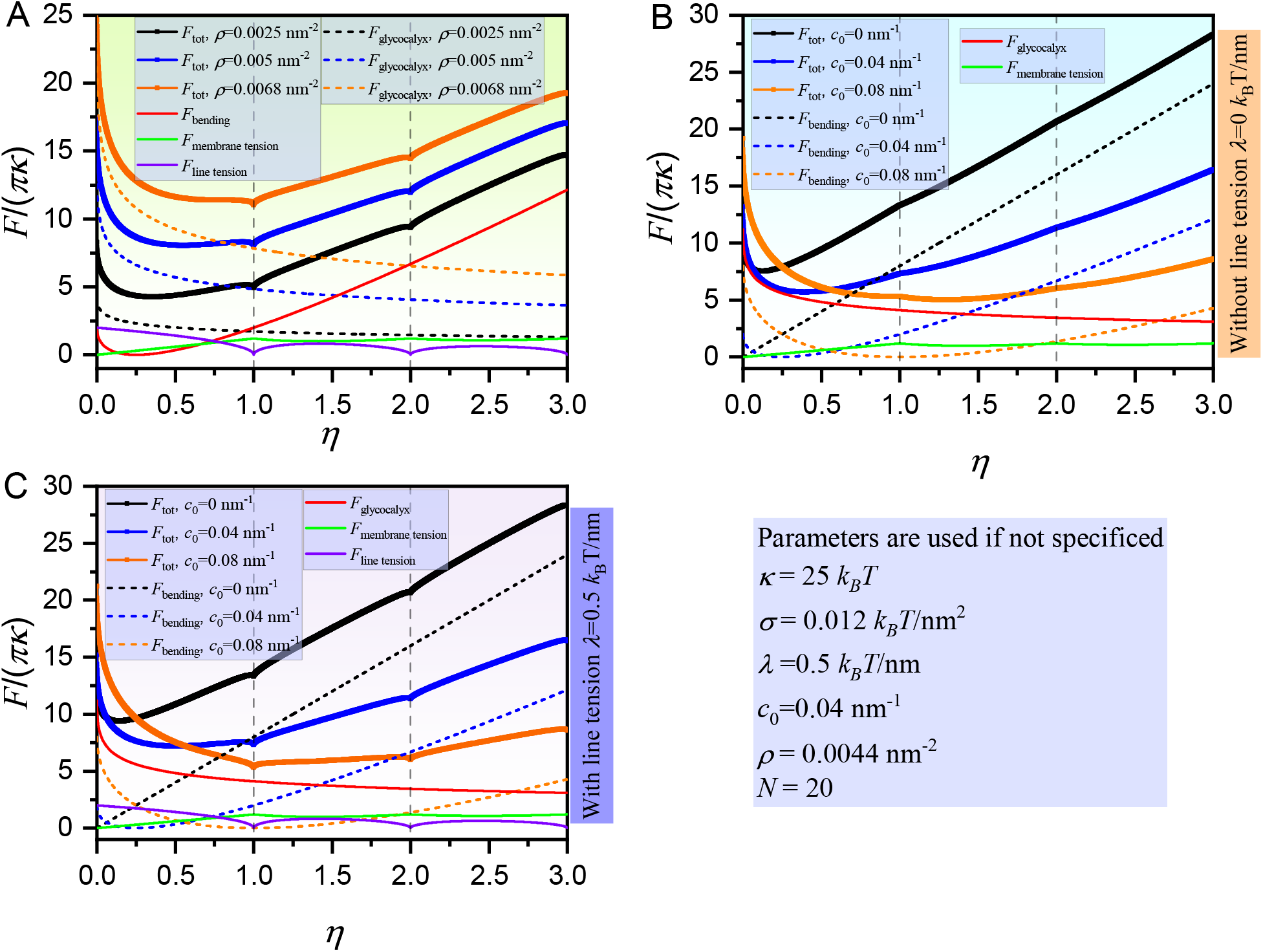
Energy contributions in membrane shape remodeling. Different types of energy profiles including total energy *F*_tot_, energy contribution associated with glycocalyx *F*_glycocalyx_, bending energy *F*_bending_, tension energy *F*_membrane tension_, and energy associated with the line tension *F*_line tension_ as a function of the shape parameter *η* for different (A) grafting density *ρ*, and spontaneous curvature *c*_0_ (B) without line tension and (C) with line tension, respectively.

### 2 Curvature Sensing Model

The curvature generating and curvature sensing capabilities of polymers are tightly interconnected. To predict the relative partitioning of glycocalyx polymers on the filopodia and on the cell surface, we developed a thermodynamic model of curvature sensing of the glycocalyx polymers. Since we have established that the glycocalyx can generate curvature, here, we model the curvature generation effects of the glycocalyx into a spontaneous curvature, *c*_eff_. Since the induced spontaneous curvature is a function of polymer area fraction and its length, we assume that the spontaneous curvature is a function of the polymer area fraction *ϕ* (ratio between the membrane area anchored by polymers and the area of the entire membrane) and the number of monomers *N*. There are many models in the literature for the curvature induced by different polymer systems on the membrane (*39, 40, 43–47, 97*). We chose the model presented in (*97*) because it incorporates the effect of *ϕ* and *N* in the spontaneous curvature

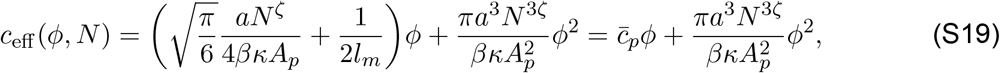

where *A*_*p*_ is the surface area occupied by the anchor, *l*_*m*_ is the membrane thickness (*l*_*m*_ ∼4− 5 nm), and the exponent *ζ*= 3*/*5 for polymer brushes under good solvent conditions. Here, for simplicity, we merely consider the linear term in Eq. (S19) at low polymer densities.

The free energy of the membrane, including the mixing energy and interaction between polymers can be written as (*56, 59–62, 98*)

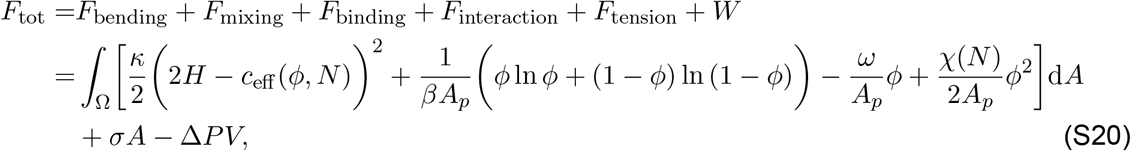

where *ω* represents the chemical potential of the polymer binding, and *χ* is a direct interaction coefficient between polymers.

To estimate the curvature sensing capability, we consider the membrane to have a spherical vesicle part representing the whole cell and a tube of radius *R*_*t*_ and length *L*_*t*_, representing the filopodia, as shown in Fig. 6A in the main text. The bending energy can then be written sum of the spherical and tubular membrane components as follows

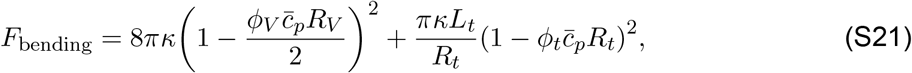

where *ϕ*_*V*_ and *ϕ*_*t*_ are the glycocalyx polymer area fractions on the sphere cell and tube, and *R*_*V*_ is the radius of the spherical cell.

Additionally, the spatial inhomogeneities in a two-component system (membrane + glycocalyx) cost mixing entropy. The second term in Eq. (S20) represents this mixing energy, which has the Flory-Huggins form, as follows

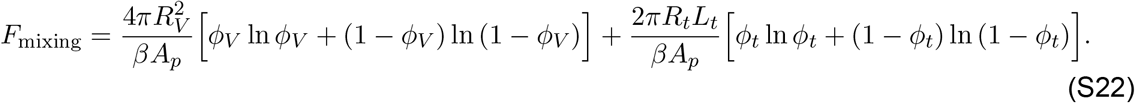

The third term is the energy associated with polymer binding to the membrane; *ω* represents the chemical potential of the polymer binding. This energy is given by

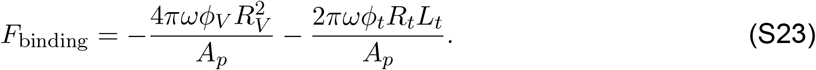

The fourth term is the energy arising from polymer-polymer interactions and is given by

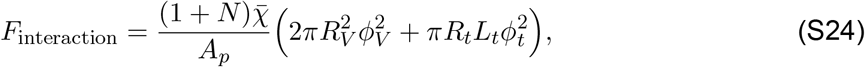

where 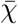 is a measure of the strength of inter-polymer interactions.

Note that the length of polymers also affects the interactions here. However, the exact dependence of *χ* on *N* is still unknown for polymers. For simplicity, we assume that they obey a linear dependence given by 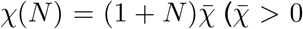 and 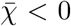 represent repulsive or attractive interactions respectively). In such a treatment, the coefficient *χ* can be reduced to the frequently encountered case that the size effect of the proteins is neglected, corresponding to 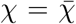 when *N* = 0.

The fifth term represents the surface tension energy and is given by

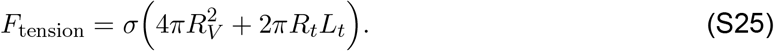

The last term is the work done by the hydrostatic pressure difference, Δ*P*, between the inside and outside of the cell

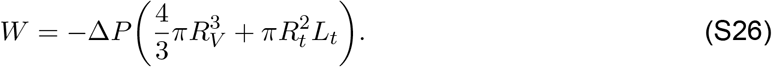

Therefore, summing Eqs. (S21)-(S26) leads to the total energy

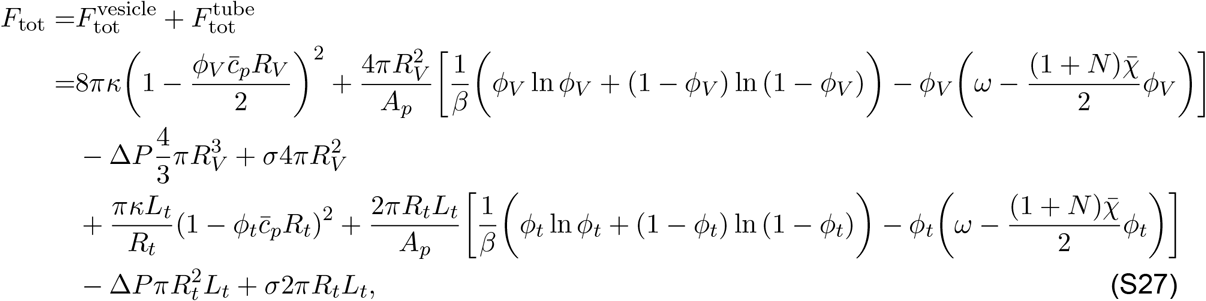

where

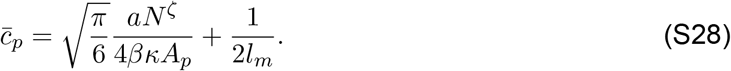

In order to quantify the curvature-sensing ability of glycocalyx, it is necessary to determine the relative partitioning of glycocalyx polymers on the filopodia and on the cell surface, which is defined by the ratio between *ϕ*_*V*_ and *ϕ*_*t*_. Therefore, we now seek to find *ϕ*_*t*_*/ϕ*_*V*_ at equilibrium. In thermal equilibrium, the polymer-bound fraction *ϕ* is locally determined for the given curvatures. By minimising the total free energy with respect to *ϕ*_*V*_, i.e., *∂F*_tot_*/∂ϕ*_*V*_ = 0, one can obtain

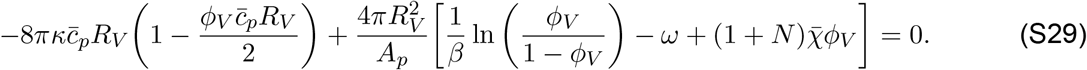

Assuming that the vesicle (and the cell) can be treated as nearly flat, 1*/R*_*V*_ = 0, the above equation reduces to

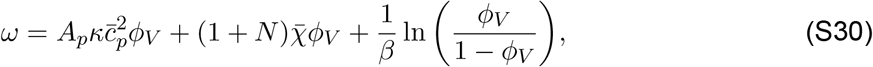

which can be solved numerically to obtain *ϕ*_*V*_. Under the special conditions that 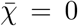 and in the absence of spontaneous curvature, we can obtain an analytical solution which denotes the equilibrium area fraction *ϕ*_V,eq_ and is obtained as a sigmoid function:

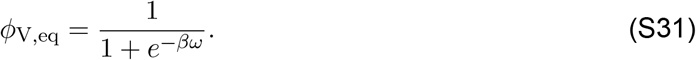

By employing perturbation theory, one can expand Eq. (S31) and keep the first order term of Δ*ϕ*_*V*_ = *ϕ*_*V*_ − *ϕ*_V,eq_, obtaining *βϕ*_V,eq_(1 − *ϕ*_V,eq_)

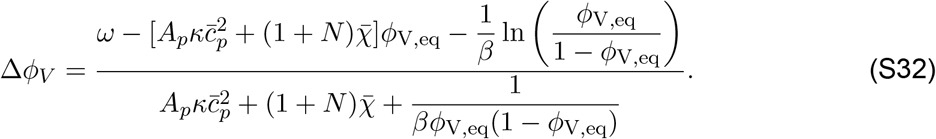

Minimization of *F*_tot_ with respect to *R*_*V*_ then leads to the Laplace law

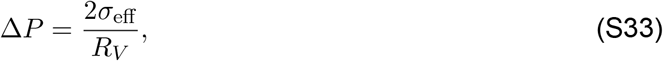

where

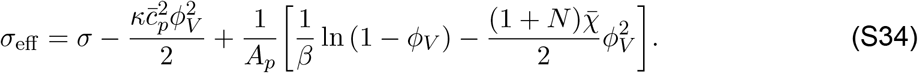

Similarly, the equilibrium fraction of polymers on the tube can be obtained by minimizing *F*_tot_ with respect to *ϕ*_*t*_, which yields

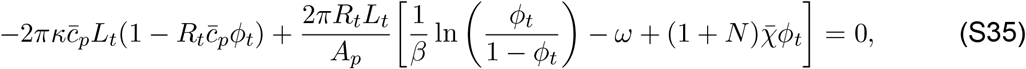

Substituting Eq. (S30) into Eq. (S35) and then expanding it to the first order in Δ*ϕ* = *ϕ*_*t*_ − *ϕ*_*V*_, one can obtain

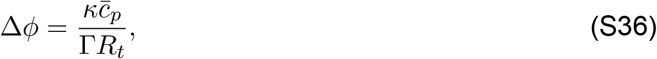

where

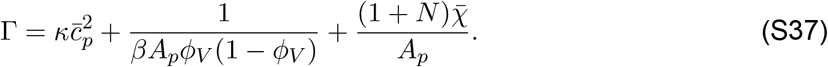

Next, minimizing *F*_tot_ with respect to *R*_*t*_ and using Eqs. (S30) and (S36) yields the tube radius

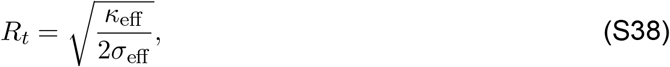

where

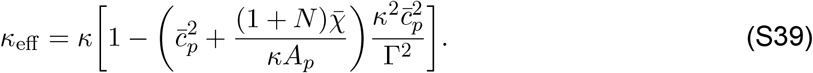

Based on these derivations, we minimize the total energy *F*_tot_ with respect to *ϕ*_*V*_, *R*_*V*_, *ϕ*_*t*_, and *R*_*t*_, respectively, to determine the relative partitioning of glycocalyx polymers on the filopodia and on the cell surface, *ϕ*_*t*_*/ϕ*_*V*_, which depends on glycopolymer length *N*. We can obtain *ϕ*_*V*_ by numerically solving Eq. S30, and then determining *ϕ*_*t*_*/ϕ*_*V*_ = 1 + Δ*ϕ/ϕ*_*V*_ by using Eq. S36. This allows us to investigate the dependence of *ϕ*_*t*_*/ϕ*_*V*_ on glycopolymer length *N*, as shown in Fig. 6 in the main text.

## Experimental materials and methods

### Cell Culture and Transfection

Human breast cancer epithelial cell line, SUM159, was cultured in 50% of Dulbecco’s modified eagle medium (Hyclone, Cytiva), 50% of F-12 nutrient mixture (Gibco), supplemented with 5% fetal bovine serum (StemCell Technologies), 1% Penicillin/Streptomycin/l-glutamine (Hyclone), 1 *µ*g*/*ml hydrocortisone (H4001; Sigma-Aldrich), 5 *µ*g*/*ml insulin (I6634; Sigma-Aldrich), and 10 mM HEPES, pH 7.4. Cells were incubated at 37 ^°^ with 5% CO_2_. SUM159 cells were seeded onto acid-washed coverslips at a density of 4*×*10^4^ cells per coverslip and incubated for 24 h. Cells were then transfected with 1 *µ*g of each plasmid using 3 *µ*g of Fugene HD (Promega). To induce expression of MUC1 ectodomain variants (TfR-GFP-0TR, TfR-GFP-10TR, MUC1-42TR-GFP), 0.05 *µ*g*/*mL of Doxycycline Hyclate (Santa Cruz Biotechnology) was added to the cell culture medium 24 h after transfection.

### Plasmids

Plasmids for the expression of TfR-GFP-0TR, TfR-GFP-10TR, and TfR-Δecto-RFP control were generated as previously described (*33, 99*). MUC1 42TR GFP ΔCT pPB was a gift from the Paszek lab (*30*) (Cornell University).

### Live cell imaging

Plasma membrane of live cells was taken by a spinning disk confocal microscope equipped with a Yokogawa CSU-W1 SoRa confocal scanner unit, an Olympus IX83 microscope body, and an Olympus 100 *×* plan-apochromat 1.5 NA oil-immersion objective. Excitation wavelengths of 488 nm and 561 nm were used for GFP and RFP, respectively.

### Image analysis

Fluorescence image analysis was conducted using FIJI software (https://imagej.net/software/fiji/). The Curvature sensing of filopodia was determined by averaging the maximum value of three perpendicular line scans. Background fluorescence was subtracted from both the membrane and filopodia intensities before determining the curvature. The Filopodyan plugin (https://github.com/gurdon-institute/Filopodyan) was employed to quantify the number of filopodia and cell perimeter.

### Statistical analysis

All live cell imaging experiments were repeated independently on different days three times, yielding consistent results. Statistical analysis was performed using one-way ANOVA with Tukey’s test for multiple comparisons within groups by GraphPad Prism 8.0. Error bars in graphs represent standard deviation.

## References

1. H. McMahon, J. Gallop, NATURE 438, 590–596, ISSN: 0028-0836, DOI 10.1038/nature04396 (Dec. 2005).

2. J. H. Hurley, E. Boura, L.-A. Carlson, B. Różycki, Cell 143, 875–887, ISSN: 0092-8674, DOI 10.1016/j.cell.2010.11.030 (2010).

3. W. F. Zeno, K. J. Day, V. D. Gordon, J. C. Stachowiak, Annual Review of Biophysics 49, PMID: 31913664, 19–39, DOI 10.1146/annurev-biophys-121219-081637, eprint: https://doi.org/10.1146/annurev-biophys-121219-081637 (2020).

4. J. C. Stachowiak, F. M. Brodsky, E. A. Miller, Nat. Cell Biol. 15, 1019–1027, ISSN: 1465-7392, DOI 10.1038/ncb2832 (Sept. 2013).

5. J. Derganc, A. čopič, Biochimica et Biophysica Acta (BBA) - Biomembranes 1858, 1152–1159, ISSN: 0005-2736, DOI 10.1016/j.bbamem.2016.03.009 (2016).

6. J. C. Stachowiak et al., NATURE CELL BIOLOGY 14, 944+, ISSN: 1465-7392, DOI 10.1038/ncb2561 (Sept. 2012).

7. D. J. Busch et al., NATURE COMMUNICATIONS 6, ISSN: 2041-1723, DOI 10.1038/ncomms8875 (July 2015).

8. S. Liese, A. Carlson, Biophysical Journal 120, 2482–2489, ISSN: 0006-3495, DOI 10.1016/j.bpj.2021.04.029 (2021).

9. V. T. Ruhoff, G. Moreno-Pescador, W. Pezeshkian, P. M. Bendix, Biochemical Society Transactions 50, 1257–1267, ISSN: 0300-5127, DOI 10.1042/BST20210883, eprint: https://portlandpress.com/biochemsoctrans/article-pdf/50/5/1257/939031/bst-2021-0883c.pdf (Oct. 2022).

10. M. M. Kozlov, J. W. Taraska, NATURE REVIEWS MOLECULAR CELL BIOLOGY 24, 63–78, ISSN: 1471-0072, DOI 10.1038/s41580-022-00511-9 (Jan. 2023).

11. H. Alimohamadi, P. Rangamani, Biomolecules 8, ISSN: 2218-273X, DOI 10.3390/biom8040120 (2018).

12. C. L. Hattrup, S. J. Gendler, Annual Review of Physiology 70, PMID: 17850209, 431–457, DOI 10.1146/annurev.physiol.70.113006.100659, eprint: https://doi.org/10.1146/annurev.physiol.70.113006.100659 (2008).

13. Y. Jung et al., Proceedings of the National Academy of Sciences 113, E5916–E5924, DOI 10.1073/pnas.1605399113, eprint: https://www.pnas.org/doi/pdf/10.1073/pnas.1605399113 (2016).

14. G. Kesavan et al., Cell 139, 791–801, ISSN: 0092-8674, DOI 10.1016/j.cell.2009.08.049 (2009).

15. M. Kesimer et al., Mucosal Immunology 6, 379–392, ISSN: 1933-0219, DOI 10.1038/mi.2012.81 (2013).

16. S. Makabe, T. Naguro, T. Stallone, Microscopy Research and Technique 69, 436–449, DOI 10.1002/jemt.20303, eprint: https://analyticalsciencejournals.onlinelibrary.wiley.com/doi/pdf/10.1002/jemt.20303 (2006).

17. S. P. Evanko, M. I. Tammi, R. H. Tammi, T. N. Wight, Advanced Drug Delivery Reviews 59, Natural and Artificial Cellular Microenvironments for Soft Tissue Repair, 1351–1365, ISSN: 0169-409X, DOI 10.1016/j.addr.2007.08.008 (2007).

18. B. Button et al., Science 337, 937–941, DOI 10.1126/science.1223012, eprint: https://www.science.org/doi/pdf/10.1126/science.1223012 (2012).

19. J. Richard Bennett et al., Journal of Histochemistry & Cytochemistry 49, PMID: 11118479, 67–77, DOI 10.1177/002215540104900107, eprint: https://doi.org/10.1177/002215540104900107 (2001).

20. V. Koistinen et al., Experimental Cell Research 337, Cell Biology at High Resolution, 179–191, ISSN: 0014-4827, DOI 10.1016/j.yexcr.2015.06.016 (2015).

21. D. W. Kufe, NATURE REVIEWS CANCER 9, 874–885, ISSN: 1474-175X, DOI 10.1038/nrc2761 (Dec. 2009).

22. E. A. Turley, D. K. Wood, J. B. McCarthy, Cancer Research 76, 2507–2512, ISSN: 0008-5472, DOI 10.1158/0008-5472.CAN-15-3114, eprint: https://aacrjournals.org/cancerres/article-pdf/76/9/2507/2748884/2507.pdf (May 2016).

23. S. Cloosen et al., International Immunology 16, 1561–1571, ISSN: 0953-8178, DOI 10.1093/intimm/dxh157, eprint: https://academic.oup.com/intimm/article-pdf/16/11/1561/2015264/dxh157.pdf (Sept. 2004).

24. L. Gangoda, S. Boukouris, M. Liem, H. Kalra, S. Mathivanan, PROTEOMICS 15, 260–271, DOI 10.1002/pmic.201400234, eprint: https://analyticalsciencejournals.onlinelibrary.wiley.com/doi/pdf/10.1002/pmic.201400234 (2015).

25. R. E. McConnell et al., Journal of Cell Biology 185, 1285–1298, ISSN: 0021-9525, DOI 10.1083/jcb.200902147, eprint: https://rupress.org/jcb/article-pdf/185/7/1285/1899464/jcb\_200902147.pdf (June 2009).

26. M. J. Paszek et al., NATURE 511, 319+, ISSN: 0028-0836, DOI 10.1038/nature13535 (July 2014).

27. T. Pelaseyed et al., Immunological Reviews 260, 8–20, DOI 10.1111/imr.12182, eprint: https://onlinelibrary.wiley.com/doi/pdf/10.1111/imr.12182 (2014).

28. J. C. Christopher Tricarico, C. D’Souza-Schorey, Small GTPases 8, PMID: 27494381, 220–232, DOI 10.1080/21541248.2016.1215283, eprint: https://doi.org/10.1080/21541248.2016.1215283 (2017).

29. R. Xu et al., NATURE REVIEWS CLINICAL ONCOLOGY 15, 617–638, ISSN: 1759-4774, DOI 10.1038/s41571-018-0036-9 (Oct. 2018).

30. C. R. Shurer et al., Cell 177, 1757–1770.e21, ISSN: 0092-8674, DOI 10.1016/j.cell.2019.04.017 (2019).

31. K. Godula, Cell 177, 1672–1674, ISSN: 0092-8674, DOI 10.1016/j.cell.2019.05.053 (2019).

32. C.-H. Lu et al., NATURE COMMUNICATIONS 13, DOI 10.1038/s41467-022-30610-2 (June 2022).

33. S. Gollapudi et al., Proceedings of the National Academy of Sciences 120, e2215815120, DOI 10.1073/pnas.2215815120, eprint: https://www.pnas.org/doi/pdf/10.1073/pnas.2215815120 (2023).

34. F. Yuan et al., Science Advances 9, eadg3485, DOI 10.1126/sciadv.adg3485, eprint: https://www.science.org/doi/pdf/10.1126/sciadv.adg3485 (2023).

35. J. R. Houser et al., Biophysical Journal 121, 3320–3333, ISSN: 0006-3495, DOI 10.1016/j.bpj.2022.08.028 (2022).

36. J. C.-H. Kuo, J. G. Gandhi, R. N. Zia, M. J. Paszek, NATURE PHYSICS 14, 658–669, ISSN: 1745-2473, DOI 10.1038/s41567-018-0186-9 (July 2018).

37. S. Weinbaum, J. M. Tarbell, E. R. Damiano, Annual Review of Biomedical Engineering 9, 121–167, ISSN: 1545-4274, DOI 10.1146/annurev.bioeng.9.060906.151959 (2007).

38. L. Möckl, Frontiers in Cell and Developmental Biology 8, ISSN: 2296-634X, DOI 10.3389/fcell.2020.00253 (2020).

39. C. Hiergeist, R. Lipowsky, JOURNAL DE PHYSIQUE II 6, 1465–1481, ISSN: 1155-4312 (Oct. 1996).

40. R. Lipowsky, Europhysics Letters 30, 197, DOI 10.1209/0295-5075/30/4/002 (May 1995).

41. M. Breidenich, R. R. Netz, R. Lipowsky, Europhysics Letters 49, 431, DOI 10.1209/epl/i2000-00167-2 (Feb. 2000).

42. T. Bickel, C. Jeppesen, C. Marques, EUROPEAN PHYSICAL JOURNAL E 4, 33–43, ISSN: 1292-8941, DOI 10.1007/s101890170140 (Jan. 2001).

43. Y. W. Kim, W. Sung, Phys. Rev. E 63, 041910, DOI 10.1103/PhysRevE.63.041910 (4 Mar. 2001).

44. F. Campelo, A. Hernández-Machado, Phys. Rev. Lett. 99, 088101, DOI 10.1103/PhysRevLett.99.088101 (8 Aug. 2007).

45. F. Campelo, A. Hernández–Machado, Phys. Rev. Lett. 100, 158103, DOI 10.1103/PhysRevLett.100.158103 (15 Apr. 2008).

46. M. Werner, J..-. Sommer, EUROPEAN PHYSICAL JOURNAL E 31, 383–392, ISSN: 1292-8941, DOI 10.1140/epje/i2010-10576-4 (Apr. 2010).

47. S. Kutti Kandy, R. Radhakrishnan, Biophysical Journal 121, 3674–3683, ISSN: 0006-3495, DOI 10.1016/j.bpj.2022.05.031 (2022).

48. W. Helfrich, ZEITSCHRIFT FUR NATURFORSCHUNG C-A JOURNAL OF BIOSCIENCES C 28, 693–703, ISSN: 0939-5075, DOI 10.1515/znc-1973-11-1209 (1973).

49. E. B. Zhulina, T. M. Birshtein, O. V. Borisov, EUROPEAN PHYSICAL JOURNAL E 20, 243–256, ISSN: 1292-8941, DOI 10.1140/epje/i2006-10013-5 (July 2006).

50. T. Baumgart, B. R. Capraro, C. Zhu, S. L. Das, Annual Review of Physical Chemistry 62, 483–506, ISSN: 1545-1593, DOI 10.1146/annurev.physchem.012809.103450 (2011).

51. B. R. Capraro, Y. Yoon, W. Cho, T. Baumgart, Journal of the American Chemical Society 132, PMID: 20050657, 1200–1201, DOI 10.1021/ja907936c, eprint: https://doi.org/10.1021/ja907936c (2010).

52. R. P. Bradley, R. Radhakrishnan, Proceedings of the National Academy of Sciences 113, E5117–E5124, DOI 10.1073/pnas.1605259113, eprint: https://www.pnas.org/doi/pdf/10.1073/pnas.1605259113 (2016).

53. C. Prevost et al., NATURE COMMUNICATIONS 6, ISSN: 2041-1723, DOI 10.1038/ncomms9529 (Oct. 2015).

54. A. Breuer, L. Lauritsen, E. Bertseva, I. Vonkova, D. Stamou, Soft Matter 15, 9829–9839, DOI 10.1039/C9SM01185D (48 2019).

55. B. Nepal, J. Leveritt, T. Lazaridis, Biophysical Journal 114, 2128–2141, ISSN: 0006-3495, DOI 10.1016/j.bpj.2018.03.030 (2018).

56. B. Sorre et al., Proceedings of the National Academy of Sciences 109, 173–178, DOI 10.1073/pnas.1103594108, eprint: https://www.pnas.org/doi/pdf/10.1073/pnas.1103594108 (2012).

57. C.-H. Lu et al., NATURE COMMUNICATIONS 13, DOI 10.1038/s41467-022-30610-2 (June 2022).

58. S. T. Milner, Science 251, 905–914, DOI 10.1126/science.251.4996.905, eprint: https://www.science.org/doi/pdf/10.1126/science.251.4996.905 (1991).

59. B. Antonny, Annual Review of Biochemistry 80, 101–123, ISSN: 1545-4509, DOI 10.1146/annurev-biochem-052809-155121 (2011).

60. P. Singh, P. Mahata, T. Baumgart, S. L. Das, Phys. Rev. E 85, 051906, DOI 10.1103/PhysRevE.85.051906 (5 May 2012).

61. Z. Shi, T. Baumgart, Nature Communications 6, ISSN: 2041-1723, DOI 10.1038/ncomms6974 (Jan. 2015).

62. T. V. S. Krishnan, S. L. Das, P. B. S. Kumar, PRAMANA-JOURNAL OF PHYSICS 94, ISSN: 0304-4289, DOI 10.1007/s12043-020-1915-z (Feb. 2020).

63. R. Lipowsky, Biophysical Journal 64, 1133–1138, ISSN: 0006-3495, DOI 10.1016/S0006-3495(93)81479-6 (1993).

64. J. Liu, M. Kaksonen, D. G. Drubin, G. Oster, Proceedings of the National Academy of Sciences 103, 10277–10282, DOI 10.1073/pnas.0601045103, eprint: https://www.pnas.org/doi/pdf/10.1073/pnas.0601045103 (2006).

65. J.-M. Allain, C. Storm, A. Roux, M. B. Amar, J.-F. Joanny, Phys. Rev. Lett. 93, 158104, DOI 10.1103/PhysRevLett.93.158104 (15 Oct. 2004).

66. L. Foret, EUROPEAN PHYSICAL JOURNAL E 37, ISSN: 1292-8941, DOI 10.1140/epje/i2014-14042-1 (May 2014).

67. Z. Shi, T. Baumgart, Nature Communications 6, ISSN: 2041-1723, DOI 10.1038/ncomms6974 (Jan. 2015).

68. R. Phillips, T. Ursell, P. Wiggins, P. Sens, NATURE 459, 379–385, ISSN: 0028-0836, DOI 10.1038/nature08147 (May 2009).

69. T. Baumgart, S. Hess, W. Webb, NATURE 425, 821–824, ISSN: 0028-0836, DOI 10.1038/nature02013 (Oct. 2003).

70. R. Lipowsky, JOURNAL DE PHYSIQUE II 2, 1825–1840, ISSN: 1155-4312 (Oct. 1992).

71. D. Bracha, E. Karzbrun, G. Shemer, P. A. Pincus, R. H. Bar-Ziv, Proceedings of the National Academy of Sciences 110, 4534–4538, DOI 10.1073/pnas.1220076110, eprint: https://www.pnas.org/doi/pdf/10.1073/pnas.1220076110 (2013).

72. L. Foret, P. Sens, Proceedings of the National Academy of Sciences 105, 14763–14768, DOI 10.1073/pnas.0801173105, eprint: https://www.pnas.org/doi/pdf/10.1073/pnas.0801173105 (2008).

73. J. Paturej, S. S. Sheiko, S. Panyukov, M. Rubinstein, Science Advances 2, e1601478, DOI 10.1126/sciadv.1601478, eprint: https://www.science.org/doi/pdf/10.1126/sciadv.1601478 (2016).

74. J. G. Gandhi, D. L. Koch, M. J. Paszek, Biophysical Journal 116, 694–708, ISSN: 0006-3495, DOI 10.1016/j.bpj.2018.12.023 (2019).

75. J. E. Hassinger, G. Oster, D. G. Drubin, P. Rangamani, Proceedings of the National Academy of Sciences 114, E1118–E1127, DOI 10.1073/pnas.1617705114, eprint: https://www.pnas.org/doi/pdf/10.1073/pnas.1617705114 (2017).

76. A. Mahapatra, P. Rangamani, Soft Matter 19, 4345–4359, DOI 10.1039/D2SM01676A (23 2023).

77. K. Xiao, R. Ma, C.-X. Wu, Phys. Rev. E 106, 044411, DOI 10.1103/PhysRevE.106.044411 (4 Oct. 2022).

78. A. Jacinto, L. Wolpert, Current Biology 11, R634, ISSN: 0960-9822, DOI 10.1016/S0960-9822(01)00378-5 (2001).

79. T. C. A. Blake, J. L. Gallop, en, Annu. Rev. Cell Dev. Biol. 39, 307–329 (Oct. 2023).

80. I. Tsafrir et al., Phys. Rev. Lett. 86, 1138–1141, DOI 10.1103/PhysRevLett.86.1138 (6 Feb. 2001).

81. I. Tsafrir, Y. Caspi, M.-A. Guedeau-Boudeville, T. Arzi, J. Stavans, Phys. Rev. Lett. 91, 138102, DOI 10.1103/PhysRevLett.91.138102 (13 Sept. 2003).

82. C. Beyer, D. S. Pisetsky, en, Nat. Rev. Rheumatol. 6, 21–29 (Jan. 2010).

83. J. Ikenouchi, K. Aoki, en, FEBS J. 289, 7907–7917 (Dec. 2022).

84. G. T. Charras, C.-K. Hu, M. Coughlin, T. J. Mitchison, en, J. Cell Biol. 175, 477–490 (Nov. 2006).

85. L. L. Norman, J. Brugués, K. Sengupta, P. Sens, H. Aranda-Espinoza, en, Biophys. J. 99, 1726–1733 (Sept. 2010).

86. I. Lavi et al., en, Biophys. J. 117, 1485–1495 (Oct. 2019).

87. R. Alert, J. Casademunt, J. Brugués, P. Sens, en, Biophys. J. 108, 1878–1886 (Apr. 2015).

88. A. Mahapatra, S. A. Malingen, P. Rangamani, en, bioRxiv, 2024.02.07.579325 (Feb. 2024).

89. H. Kang et al., International Journal of Molecular Sciences 19, ISSN: 1422-0067, DOI 10.3390/ijms19092484 (2018).

90. N. Charnaux et al., en, Glycobiology 15, 119–130 (Feb. 2005).

91. R. Uchimido, E. P. Schmidt, N. I. Shapiro, en, Crit. Care 23, 16 (Jan. 2019).

92. B. F. Becker, M. Jacob, S. Leipert, A. H. J. Salmon, D. Chappell, en, Br. J. Clin. Pharmacol. 80, 389–402 (Sept. 2015).

93. T. M. Handel, D. P. Dyer, en, J. Histochem. Cytochem. 69, 87–91 (Feb. 2021).

94. C. Zhu, C. T. Lee, P. Rangamani, Biophysical Reports 2, 100062, ISSN: 2667-0747, DOI 10.1016/j.bpr.2022.100062 (2022).

95. C. T. Lee, P. Rangamani, in Biophysical Approaches for the Study of Membrane Structure—Part B: Theory and Simulations, ed. by M. Deserno, T. Baumgart (Academic Press, 2024), vol. 701, pp. 387–424, DOI 10.1016/bs.mie.2024.03.016.

96. T. Baumgart, S. Hess, W. Webb, NATURE 425, 821–824, ISSN: 0028-0836, DOI 10.1038/nature02013 (Oct. 2003).

97. V. Nikolov, R. Lipowsky, R. Dimova, Biophysical Journal 92, 4356–4368, ISSN: 0006-3495, DOI 10.1529/biophysj.106.100032 (2007).

98. N. S. Gov, Philosophical Transactions of the Royal Society B: Biological Sciences 373, 20170115, DOI 10.1098/rstb.2017.0115, eprint: https://royalsocietypublishing.org/doi/pdf/10.1098/rstb.2017.0115 (2018).

99. W. F. Zeno et al., NATURE COMMUNICATIONS 9, ISSN: 2041-1723, DOI 10.1038/s41467-018-06532-3 (Oct. 2018).

